# Interplay of EGFR, JNK and ROS signalling in soma-germline communication in the *Drosophila* testis

**DOI:** 10.1101/2024.06.02.597033

**Authors:** Maria Alvarez, Fani Papagiannouli

## Abstract

Cell communication via signalling exchange plays a pivotal role in multicellular development for building functional tissues and organs. In the *Drosophila* testis, a pair of somatic cyst cells (CCs) encapsulate the germline that differentiates through close-range EGFR signalling activation. The conserved Dlg/Scrib/Lgl cortical polarity complex and clathrin-mediated endocytosis attenuate EGFR signalling in CCs and loss of their function leads to EGFR overactivation and non-autonomous death of the neighbouring germ cells. Here we show that EGFR overactivation results in upregulation of JNK and p38 signalling in CCs and ROS levels in the germ cells that are destined to die. Our data uncover a bidirectional feedback between JNK signalling and ROS who regulate each other within the CC-germline microenvironment, while reducing the levels of either JNK or ROS restores germ cell survival. This study provides a framework of how polarity and cellular trafficking regulate the output of multiple signalling responses cell-intrinsically and in adjacent cells, to coordinate tissue-specific responses and maintain homeostasis.

## Introduction

Cell-to-cell communication and exchange of short-range signals is crucial for cells to communicate with their neighbours, coordinate their function to the local microenvironment and build functional tissues and organs. Based on the signals they receive, cells adapt their intrinsic features and signalling machinery to achieve a coordinated output. In the *Drosophila* testis, cell communication via close-range signalling exchange is established between the germline and the somatic lineage that is crucial to build a functional testis and produce fertile sperm (Papagiannouli and Lohmann, 2012; Papagiannouli and Lohmann, 2015). At the anterior-most part of the testis is the male stem cell niche, a cluster of non-dividing tightly packed somatic cells that build the so-called hub. Around the hub are organised the germline stem cells (GSCs), each surrounded by a pair of the somatic cyst stem cells (CySCs) (Fig.1A). Upon asymmetric cell division, each GSC produces a new GSC attached to the hub and a daughter gonialblast (GB) that becomes displaced from the hub and enters the differentiation program (Fuller and Spradling, 2007). Along this differentiation process, the gonialblast enters a stage of 4 transit-amplifying (TA) mitotic divisions with incomplete cytokinesis, giving rise to 2-, 4-, 8- and 16-interconnected spermatogonial germ cells. Immediately after reaching the 16-cell stage, germ cells undergo a final round of DNA synthesis, enter meiotic prophase, and turn on the spermatocyte transcription program for meiosis and spermatid differentiation. CySCs also divide asymmetrically to renew themselves and produce two distally located daughter somatic cyst cells (CCs) that cease mitotic divisions and encapsulate the germline from initial differentiation to mature sperm production (Leatherman, 2013; Zoller and Schulz, 2012). As CCs encapsulate the germ cells they build together a testicular cyst, an organoid-like differentiation unit, which is further reinforced by the establishment of an isolating permeability barrier by the septate junctions (SJs) (Fairchild et al., 2015). The permeability barrier is normally established at 4-cell stage spermatogonial cysts at the contact sides of the two CCs comprising the cyst and becomes fully functional at spermatocyte-stage (Brantley and Fuller, 2019; Fairchild et al., 2015; Papagiannouli et al., 2019).

**Figure 1:**
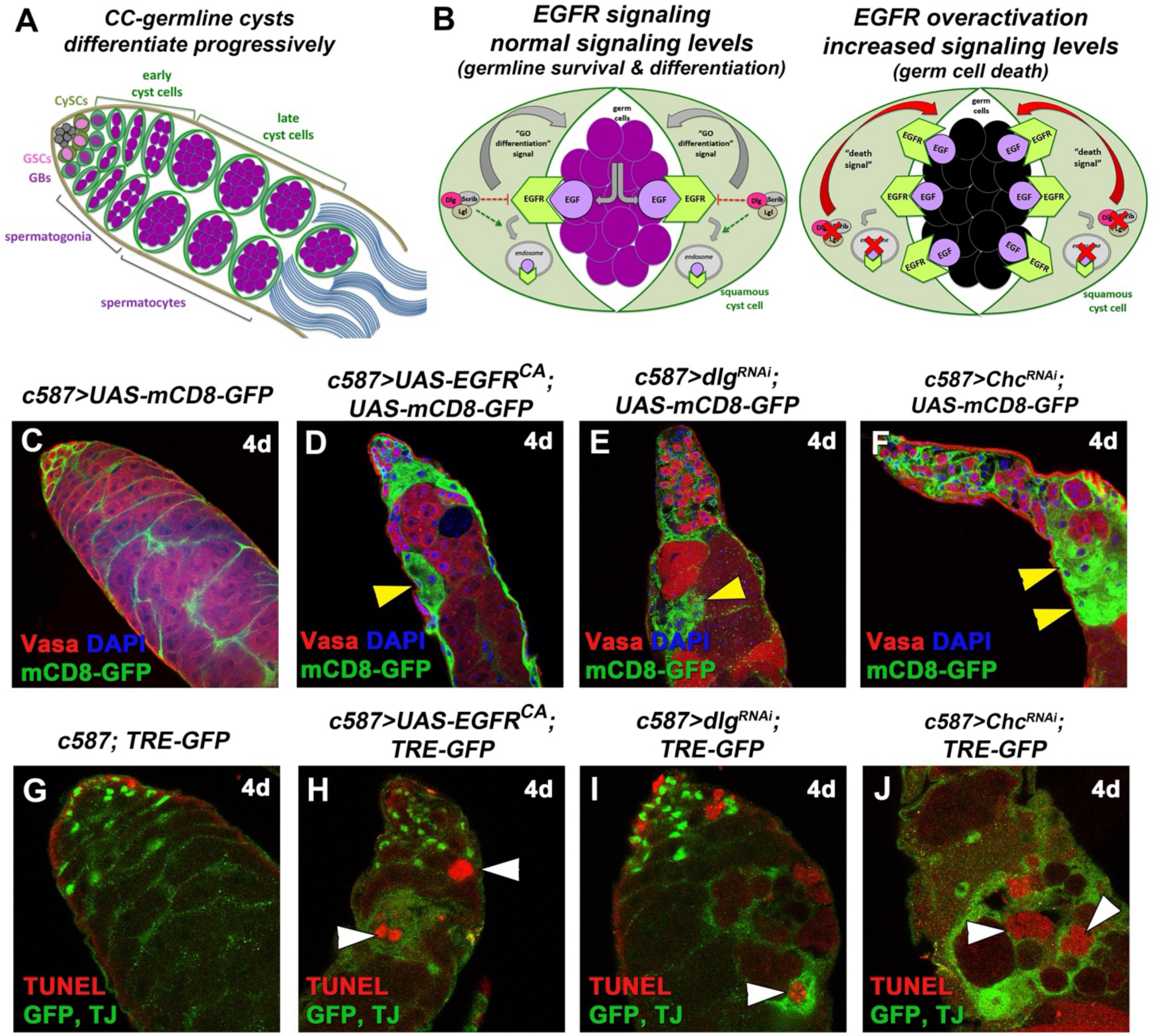
Overactivation of EGFR, also via knockdown of *dlg* or *Chc* function, in cyst cells leads to activation of JNK signalling in cyst cells and apoptosis in the neighbouring germline. **(A)** Diagram of early spermatogenesis in *Drosophila*. GSC: germline stem cell, GB: gonialblast, CySC: somatic cyst stem cell. **(B)** Diagram depicting the role of Dlg-module and CME components in EGFR signalling regulation. Upon binding the EGF-ligand Spitz from the germline, activated EGFR sends a (unidentified) “GO differentiation” signal that promotes progressive germline differentiation, while Dlg/Scrib/Lgl and CME fine-trim EGFR signalling levels. Loss of Dlg-module or CME components, leads to EGFR overactivation in CCs that sends now a “Death signal” to the germ cells (black). **(C-J)** Adult testes of the indicated genotypes in the *Gal80^ts^*background. **(C-F)** *mCD8-GFP* (green; CCs), Vasa (red; germline), DAPI (blue; nuclei). Yellow arrowheads: mCD8+ CCs regions. **(G-J)** TUNEL (red; apoptotic double-strand breaks), AP-1 responsive TRE elements corresponding to JNK reporter *puc* expression levels (*TRE-GFP*) and TJ (early CC nuclei) (green). *UAS* activated at 30°C for 4 days (4d). White arrowheads: dying germ cells (spermatogonia and spermatocytes) surrounded by CCs with upregulated JNK levels (green). Testes oriented with anterior at left. Image frame (C-F) 225μm and (G-J) 112,5μm

CCs support the progressive steps of germ cell differentiation through increasing levels of the Epidermal Growth Factor (EGFR) signalling activity (Fig.1B) (Hudson et al., 2013; Kiger et al., 2000; Sarkar et al., 2007; Tran et al., 2000). Signals from the germline to the CCs via the EGF ligand Spitz activate the EGFR in CCs and the downstream Ras/MAPK signal transduction pathway, that leads to the double phosphorylated(dp)ERK entering the nucleus (Hudson et al., 2013; Kiger et al., 2000; Sarkar et al., 2007; Tran et al., 2000). This first step of differentiation is required for germ cells to execute the TA mitotic divisions, while higher levels of EGFR activation are required for germ cells to exit TA divisions and initiate the pre-meiotic program (Hudson et al., 2013). In the absence of EGFR-derived signals, germ cells cannot differentiate and overproliferate as stem cell-like germ cells (Kiger et al., 2000; Tran et al., 2000), while overexpression of EGFR induces germ cell death (GCD) (Papagiannouli et al., 2019), highlighting the importance of fine-tuning signalling levels for a coordinated response.

In our previous work we have shown the cortical polarity proteins Discs large (Dlg), Scribble (Scrib), Lethal (2) giant larvae (Lgl) and clathrin-mediated endocytosis (CME) components attenuate EGFR signalling in CCs to promote germ cell survival and differentiation in adult male testes (Papagiannouli, 2022; Papagiannouli et al., 2019). Dlg, Scrib, Lgl, collectively called the Dlg-module, are highly conserved proteins, involved in establishment and maintenance of apical/basal polarity, signalling regulation, vesicle and membrane trafficking (Gaudet et al., 2011; Gui et al., 2016; Lohia et al., 2012; Nagasaka et al., 2013; Nakajima, 2021; Papagiannouli and Mechler, 2012; Parsons et al., 2014; Stephens et al., 2018). They are required in the somatic lineage of embryonic gonads and larval testes for the differentiation and survival of the developing *Drosophila* testis (Marhold et al., 2003; Papagiannouli, 2013; Papagiannouli and Mechler, 2009; Papagiannouli and Mechler, 2010), in a function that is non-tumorigenic, unlike epithelia in other tissues (Bilder, 2001; Bilder et al., 2000; Donohoe et al., 2018; Humbert, 2015; Stephens et al., 2018; Uhlirova and Bohmann, 2006). On the other hand, Clathrin heavy chain (Chc), Shibire (Shi; the *Drosophila* homologue of Dynamin) and AP-2α (also known as α-Adaptin) are core components of CME, which have been previously involved in EGFR signalling regulation by removing active EGFR receptor molecules from the cell surface membranes in many tissues, while recycling of EGFR back to the membrane via the action of the Rab11-recycling endosome keeps the signalling active and in physiological equilibrium (Conte and Sigismund, 2016; Dobrowolski and De Robertis, 2011; Papagiannouli, 2022; Welz et al., 2014). Knockdown of any of the Dlg-module or CME components in adult testis CCs increases EGFR signalling levels, resulting in cell non-autonomous GCD of both the spermatogonia and spermatocytes they encapsulate. Interestingly, lowering the levels of EGFR signal transduction components can rescue the observed defects and restore germ cell survival (Papagiannouli et al., 2019). Yet, the “death” signal that CCs send to the differentiating germ cells (Fig.1B), as a result of EGFR overactivation is currently unknown.

Reactive Oxygen Species (ROS) are oxygen derivatives produced as a result of normal aerobic metabolism. In physiological levels, ROS act as secondary signalling molecules in cell biology and redox signalling (Lennicke and Cocheme, 2021; Schieber and Chandel, 2014; Sinenko et al., 2021). Yet, elevated ROS levels can induce oxidative modifications resulting in cellular damage and cell death termed “oxidative stress”. ROS can activate redox-sensitive signals and the mitogen-activated protein kinases (MAPK) of the Jun N-terminal kinase (JNK) and p38 MAPK signalling pathways, through the MAPKKK Apoptotic signal-regulating kinase 1 (Ask1) that is particularly sensitive to ROS oxidative stress (Patel et al., 2019; Sakauchi et al., 2017; Santabarbara-Ruiz et al., 2019; Serras, 2022). JNK and p38 are conserved stress signalling pathways that can activate apoptosis as one of their several context-dependent and cell-specific functions (Herrera et al., 2021; La Marca and Richardson, 2020; Serras, 2022; Wagner and Nebreda, 2009). JNK signalling initiates through the activation of either of the two tumour necrosis factor (TNF) receptors Wengen (Wgn) and Grindelwald (Grnd), followed by a series of phosphorylation events leading to phosphorylation of Basket (Bsk), the sole *Drosophila* JUN kinase homologue. In turn, Bsk phosphorylates Kayak and Jun-related antigen (Jrn), the *Drosophila* homologues of Fos and Jun, respectively, which build together the AP-1 transcription factor that activates expression of downstream genes such as *puckered (puc)* and *Matrix metalloprotease (Mmp1)* (Martin-Blanco et al., 1998; Uhlirova and Bohmann, 2006). p38 is downstream of a phosphorylation cascade in which Ask1 phosphorylates the serine/threonine kinase Licorne (Lic), which can activate p38a and p38b but not p38c in *Drosophila*. Phosphorylated(p)-p38 can then activate Duox, an NAD(P)H oxidase that promotes ROS activation (Patel et al., 2019; Santabarbara-Ruiz et al., 2019; Takeda et al., 2008).

In this study, we set to explore the signal that mediates germ cell death (GCD) upon EGFR upregulation in *Drosophila* testis CCs. Our data showed that EGFR overactivation in CCs leads to increased levels of JNK and p38 signalling in CCs as well as an increase in ROS oxidative stress levels in germ cells that are destined to die. Reducing JNK levels by knocking down the JUN-kinase *bsk* in CCs, normalized p38 levels, reduced the levels of ROS in the germ cells and reversed GCD. Conversely, reducing ROS levels by feeding the flies with antioxidant Vitamin C, restored germ cell survival as well as JNK signalling in CCs to physiological levels. Our data establish a link between JNK and ROS signalling in the *Drosophila* testis that is coupled to a bidirectional feedback mechanism between the CCs and the differentiating (spermatogonia and spermatocyte) germ cells, also mediated by the JNK receptor Wengen, the p38 MAPK signalling and the recycling endosome GTPase Rab35.

## Results

### Overactivation of EGFR, also via knockdown of Dlg-module or CME components, in cyst cells leads to non-autonomous death of the neighbouring differentiating germ cells

In the adult *Drosophila* testis, upon asymmetric GSC division the (daughter) gonialblast becomes displaced from the hub and enters a differentiation process that consists of 4-rounds of TA mitotic divisions as spermatogonia, followed by a premeiotic gene expression stage as spermatocytes, meiosis and terminal differentiation leading the sperm formation. Here, the densely packed spermatogonia and spermatocytes inside the testis were marked with Vasa, while the postmitotic cyst cells (CCs) encapsulating them were marked with a membrane(m)CD8-GFP (*UAS-mCD8-GFP*) under the control of the CC lineage *c587-GAL4* driver (Fig.1C, S1A). The nuclei of early CCs (encapsulating primarily spermatogonia) were stained for the Traffic-Jam (TJ) transcription factor (Fig.1G, S1F). The function of the EGFR, Dlg-module and CME components was impaired in CCs using the *c587-GAL4* driver together with *UAS* transgenes to overexpress a constitutively active EGFR (*EGFR^CA^*) or to knockdown the expression of *dlg, scrib, lgl, Chc, shi or AP-2α* (Fig.1D-F, 1H-J, S1B-E, S1G-J). Flies also carried an *αtubGal80^ts^*transgene, which blocks *GAL4* activity at 18°C but activates *GAL4* activity at 30°C, allowing overexpression or downregulation of gene function in a timely controlled manner, after normal testis anatomy had been set up.

Analysis of the tissue 4 and 7 days after the shift to 30°C allowed us to observe the progression of the phenotypes over time. Similar to what we have previously shown (Papagiannouli et al., 2019), EGFR overactivation in CCs for 4 days (either by forced expression of *EGFR^CA^* or knockdown of Dlg- or CME-module components, thereafter called “EGFR overactivation”), led to loss of spermatogonia and spermatocytes via apoptosis, visualized by visible gaps in the Vasa staining (Fig.1D-F, S1B-E). TUNEL staining marked the dying germ cells, which appeared as red-blebs devoid of cell-type specific markers, at more advanced stages of apoptosis (white arrowheads in Fig.1H-J, S1G-J). CCs clustered together creating large “patches” marked by mCD8+ areas, in the absence of the germ cells they normally encapsulate (yellow arrowheads in Fig.1D-F, S1B-E). However, the underlying signal that mediates germ cell death (GCD) upon EGFR overactivation in CCs remained so far uncharacterized.

### Overactivation of EGFR in cyst cells leads to upregulation of JNK signalling in cyst cells and of ROS oxidative stress in the neighbouring germline

The Jun N-terminal kinase (JNK) pathway is an evolutionary conserved kinase cascade with an important role in stress-induced apoptosis and tumour progression (Herrera et al., 2021). In normal, physiological conditions, JNK signalling is activated in CySC and CCs via the JUN kinase Basket (Bsk), leading to the activation of the downstream effectors Puckered (Puc) and Matrix metalloprotease (Mmp1) (Herrera et al., 2021). Here, basal JNK levels were observed in control testes by monitoring protein levels of Mmp1 (Fig. S1K, S1M) and AP-1 responsive TRE elements fused to GFP (*puc::TRE-GFP*) (Chatterjee and Bohmann, 2012), with the latter reflecting *puc* expression levels in CCs (Fig.1G, 2A, 2E, S1F, S2A, S2F, S2K). Overexpression of *EGFR^CA^*, showed Mmp1 staining in CCs that clustered together after 4- and 7-days of activation (Fig.S1L, S1N; yellow arrowheads). Quantification of corrected Mmp1 fluorescence confirmed the increased Mmp1 levels for both 4- and 7-days (**p<0.01 and ****p<0.0001, respectively), providing the first link of JNK upregulation following EGFR overactivation in CCs. Yet, as the Mmp1 staining is challenging in the *Drosophila* testis, we focused our further analysis on *puc* expression, using *puc::TRE-GFP* as a readout for JNK signalling levels.

Overexpression of *EGFR^CA^* or knockdown of Dlg-module or CME components using the *c587-GAL4*, resulted in bright GFP staining reflecting *puc* expression in clusters of CCs that encapsulate TUNEL-positive germ cells (Fig. 1H-J, S1G-J), thereby linking JNK upregulation in CCs to the death of the germ cells they encapsulate. In order to confirm JNK upregulation in this context, we looked closer at *puc* expression upon EGFR overactivation in CCs (Fig. 2A-H, S2A-O). Quantification of corrected fluorescent *puc::TRE-GFP* levels after 4-days of *UAS* activation (4d; Fig.2M), showed significant upregulation of JNK signalling levels. Each individual sample compared to the control, showed a value of ****p<0.0001 using the non-parametric two-sampled Mann–Whitney test. Similar results were also obtained when we quantified corrected fluorescent *puc::TRE-GFP* levels after 7-days (7d; Fig.S2K-O, S2P) of *EGFR^CA^* overexpression or *dlg, scrib, Chc*, *shi, AP-2α* knockdowns in CCs. More precisely, *EGFR^CA^* overexpression and *dlg* knockdown showed a stable 3-fold and 2-fold increase in 4- and 7-days activation, respectively. CME components showed 4-6-fold increase in 4 days activation, after which levels dropped to a 2.5-3.5-fold increase (****p<0.0001) compared to the control. Loss of *scrib* and *lgl* in CCs was accompanied by a 2-fold increase of *puc* levels after 4-days (****p<0.0001), which dropped to 1.5 (**p<0.01) for *scrib* and to non-significant levels for *lgl* after 7 days. The weaker phenotype observed upon loss of *lgl* was consistent with the weaker efficiency of the *UAS-lgl-RNAi* fly lines, also confirmed in our previous study (Papagiannouli et al., 2019). Importantly, knocking down *bsk* in CCs led to efficient reduction of *puc* expression levels (****p<0.0001), thereby confirming the effectiveness of the *UAS*-*bsk-RNAi* transgene in downregulating JNK signalling in these cells (Fig.2M; column 2 vs 1).

**Figure 2:**
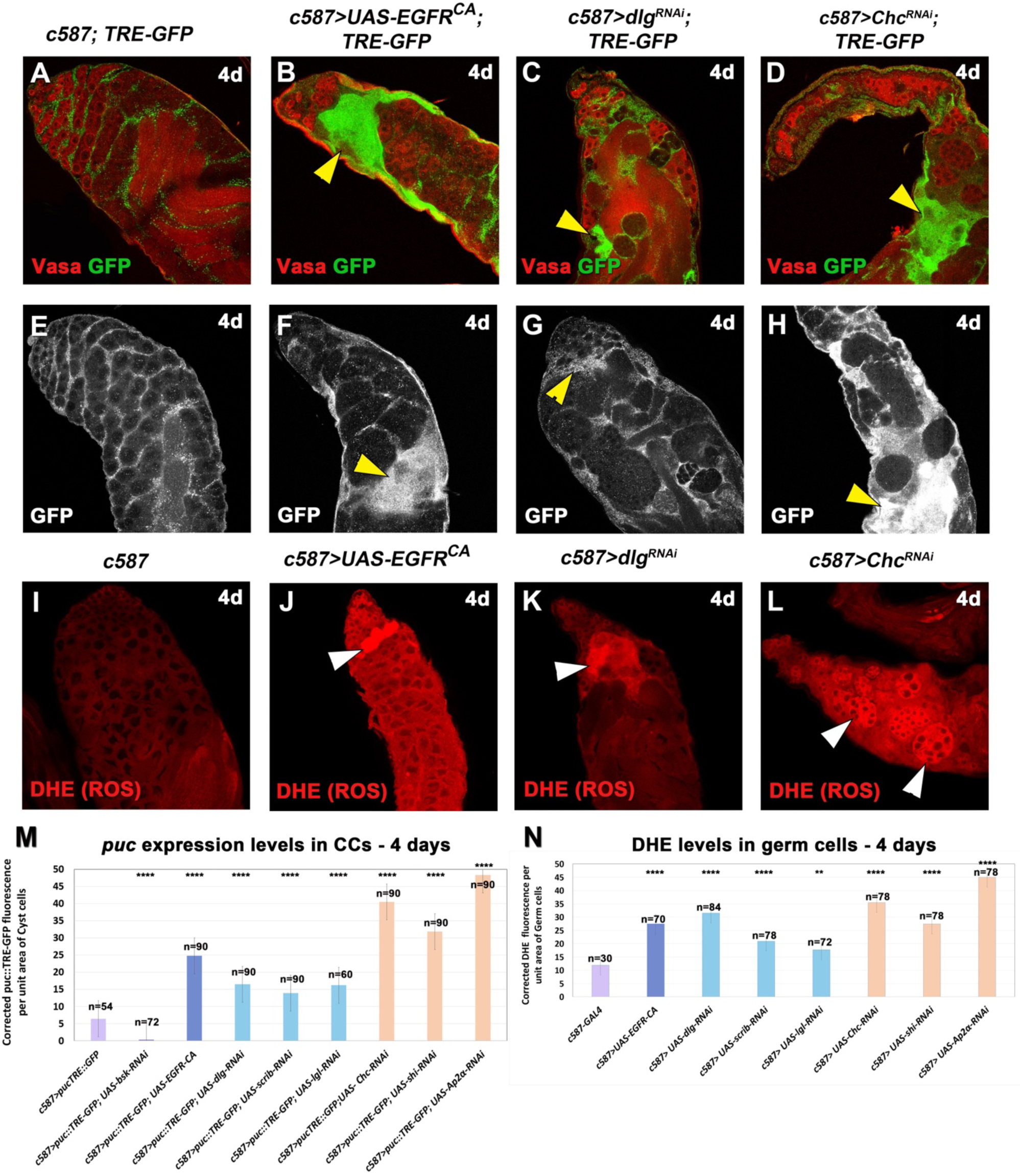
Overexpression of EGFR or knockdown of *dlg* and *Chc* function in cyst cells leads to increased levels of JNK signalling in the cyst cells and ROS in the germline. Adult testes of the indicated genotypes in the *Gal80^ts^*background. **(A-H)** *TRE-GFP* reflects expression levels of JNK reporter *puc* in CCs. (A-D) *TRE-GFP* (green; CCs); Vasa (red; germline). (E-H) show the *TRE-GFP* levels only (white) in directly comparable raw images. Yellow arrowheads: regions of *puc::TRE* overactivation in CCs. **(I-L)** DHE (red; germ cells) reflects ROS activation in the germ cells. White arrowheads: representative areas of ROS activation in the germline. *UAS* activated at 30°C for 4 days (4d). **(M, N)** Quantification of corrected fluorescent *puc::TRE-GFP* levels in CCs and DHE levels in germ cells, respectively. Each individual sample was compared to control (error bars: standard error; ns: not significant; *p<0.05; **p<0.01; ***p<0.001; ****p<0.0001). Numbers (n) in each column represent sample size. Image frames (A-L): 225μm

Reactive Oxygen Species (ROS) are oxygen derivatives acting as signalling molecules in physiological levels to promote tissue regeneration or wound healing, whereas elevated levels can induce oxidative stress resulting in cellular damage and cell death (Lennicke and Cocheme, 2021; Schieber and Chandel, 2014; Sinenko et al., 2021). Staining of testes with Dihydroethidium (DHE) to detect ROS levels, showed increased levels of ROS in differentiating germ cells upon *EGFR^CA^* overexpression or knockdown of Dlg-module and CME components in CCs (white arrowheads; Fig.2J-L, S2R-U), compared to the physiological levels of the control (Fig. 2I, S2Q). Quantification the DHE levels for each genotype after 4 days of activation (Fig.2N), confirmed these observations. All genotypes showed significant upregulation of ROS levels (****p<0.0001), while *lgl* knockdown showed significant but lower ROS activation (**p<0.01). This latter observation is most likely again due to the weaker efficiency of the *UAS-lgl-RNAi* fly lines (Papagiannouli et al., 2019).

Taken together, our results showed that EGFR overactivation in CCs (upon either forced expression of *EGFR^CA^* or loss of function of Dlg-module or CME components), led to upregulation of JNK signalling in CCs and ROS in the neighbouring germ cells prior to their death.

### Lowering the levels of the JUN kinase *basket* in cyst cells showed a partial rescue of germ cell death and reduction in ROS levels in the germline

Given the enhanced JNK activity following EGFR upregulation in CCs, we tested if reducing JNK activation might rescue the GCD phenotype. In order to interfere with JNK signalling, we used a transgenic RNAi to knockdown *bsk* in the CCs, as this was shown to effectively downregulate *puc* expression in CCs (“*bsk*-rescue”) (Fig.2M; column 2). To control for possible effects of multiple *UAS* constructs limiting the effectiveness of the *GAL4* driver, control flies were set up to carry the same number of *UAS* transgenes using a *UAS-mCD8-GFP* (we could not use a *UAS-GFP-RNAi* line, as this would interfere with the *puc::TRE-GFP*). Moreover, because different *UAS* transgenes have different expression strengths, and the phenotype manifested is the result of the equilibrium of 2 *UAS* lines, rescue experiments were performed by shifting flies to 30°C for 4 as well as 7 days.

Representative examples of the different phenotypic classes, reflecting the variability in strength and penetrance of the rescued phenotypes in comparison to control flies (wild-type-like *c587> UAS-mCD8-GFP*) are shown in Fig.3 and Fig.S3. The observed phenotypes were classified into the following categories: 1) “no rescue” for testes that mimicked the effect of acute *EGFR^CA^*overexpression in CCs [Fig.3B-D, S3B-E (4-days) and Fig.3J-L, S3L-O (7-days)] and showed decreased number of germ cells with CCs clustering together in patches devoid of germ cells. Phenotypes here ranged from milder to stronger ones; 2) “partial rescue” for testes with differentiating germ cells densely pack (and small germ cell loss if any), where encapsulation by CCs was largely restored but architecture and overall morphology appeared distorted or few CC-clusters could still be observed (Fig.3H, 3N-P, S3J, S3Q, S3S-T). 3) “Rescued (wild-type-like)” for testes with restored spermatogonial and spermatocyte cysts similar to wild type (wt) (Fig.3F-G, S3G-I, S3R).

**Figure 3.**
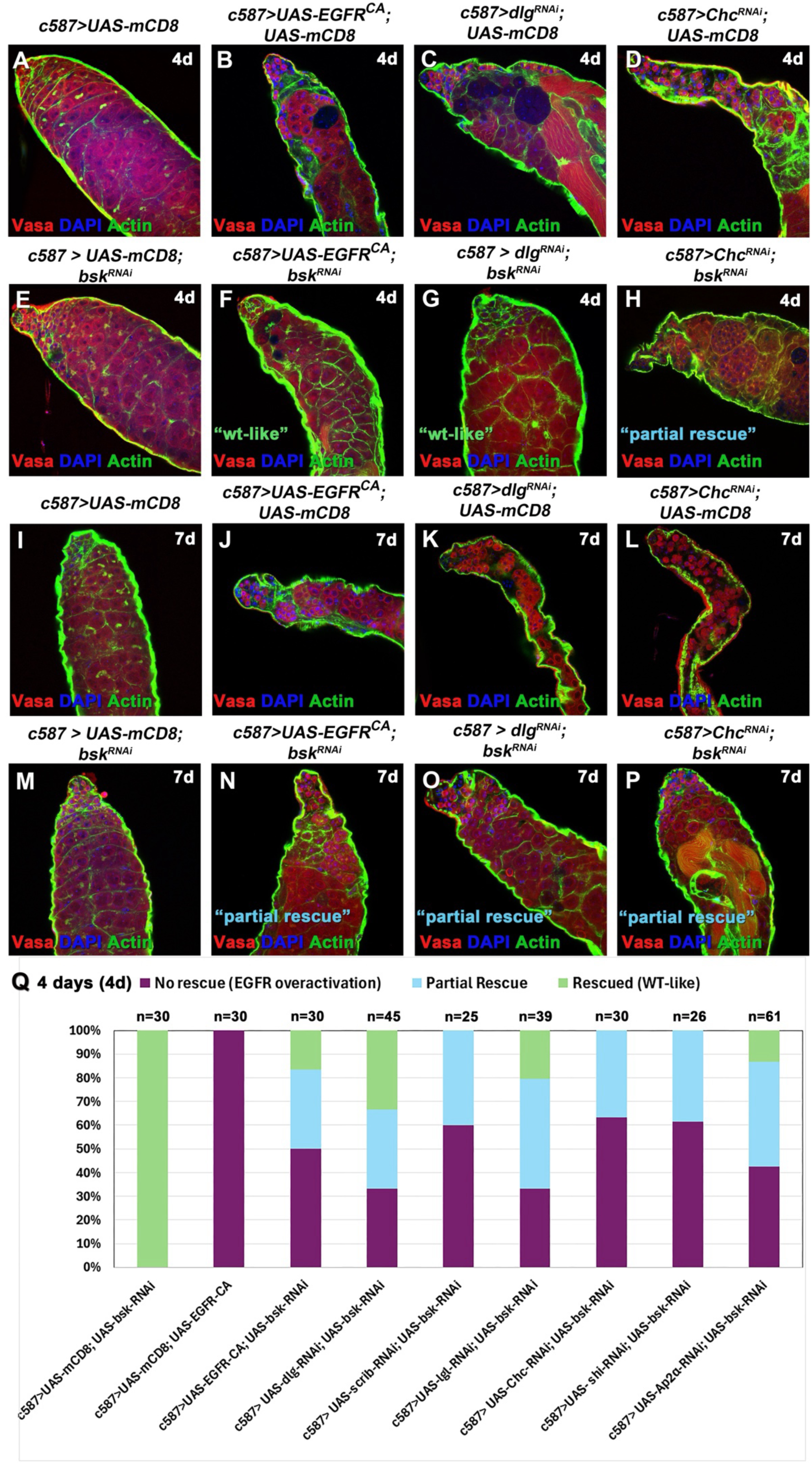
Knocking down the JUN kinase *bsk* in cyst cells, can partially rescue the germ cell death phenotype observed upon EGFR overactivation. **(A-P)** Adult testes of the indicated genotypes in the *Gal80^ts^* background: Vasa (red; germline), DAPI (blue; nuclei) and Actin stained with phalloidin (here shown in the green channel; hub, CySCs, CCs and germline fusome) also in flies containing the *mCD8-GFP* transgene (since the GFP is not shown here). *UAS* activated at 30°C for 4 and 7 days (d). **(Q)** Quantifications of the different phenotypic classes accompanying each genotype, organized in order of phenotypic strength (4d: 4 days activation). Numbers (n) in each column represent sample size. Testes oriented with anterior at left. Image frames (A-P) 225μm.

Although a range of phenotypes resulted from our rescue strategy, the percent of testes showing “no rescue” (mimicking the “EGFR overactivation” phenotype), was substantially reduced and represented 30%-62% of the overall testes scored after 4 days rescue (Fig.3Q) and 0%-63% after 7-days rescue (Fig.S3U), compared to 100% penetrance before the rescue (Fig.3Q, S3U; column 2 in both) (Papagiannouli et al., 2019). Conversely, the testes scored for full and partial rescue combined, represented a minimum of ∼40% after 4-days and minimum of 53% after 7-days, evidenced by a largely restored morphology with packed spermatogonia and spermatocyte cysts.

Effective reduction of JNK signalling in these *bsk*-rescue experiments (i.e. after knockdown of *bsk* in CCs with overactivated EGFR signalling), was confirmed by quantifying corrected *puc::TRE-GFP* fluorescence for each genotype for 4 days activation (Fig.4A-J). The results showed significant reduction in GFP fluorescence reflecting *puc* expression levels in *bsk*-rescued testes (Fig.4P), compared to the EGFR overactivation phenotypes (Fig.2M). More precisely, we could observe a significant reduction in *puc* levels upon single knockdown of *bsk* in CCs compared to the control (Fig.4P; column 2 vs 1; ****p<0.0001), and a milder reduction of *puc* levels in the *bsk*-rescue experiments (Fig.4P: columns 3-5 vs column 1; ****p<0.0001).

**Figure 4:**
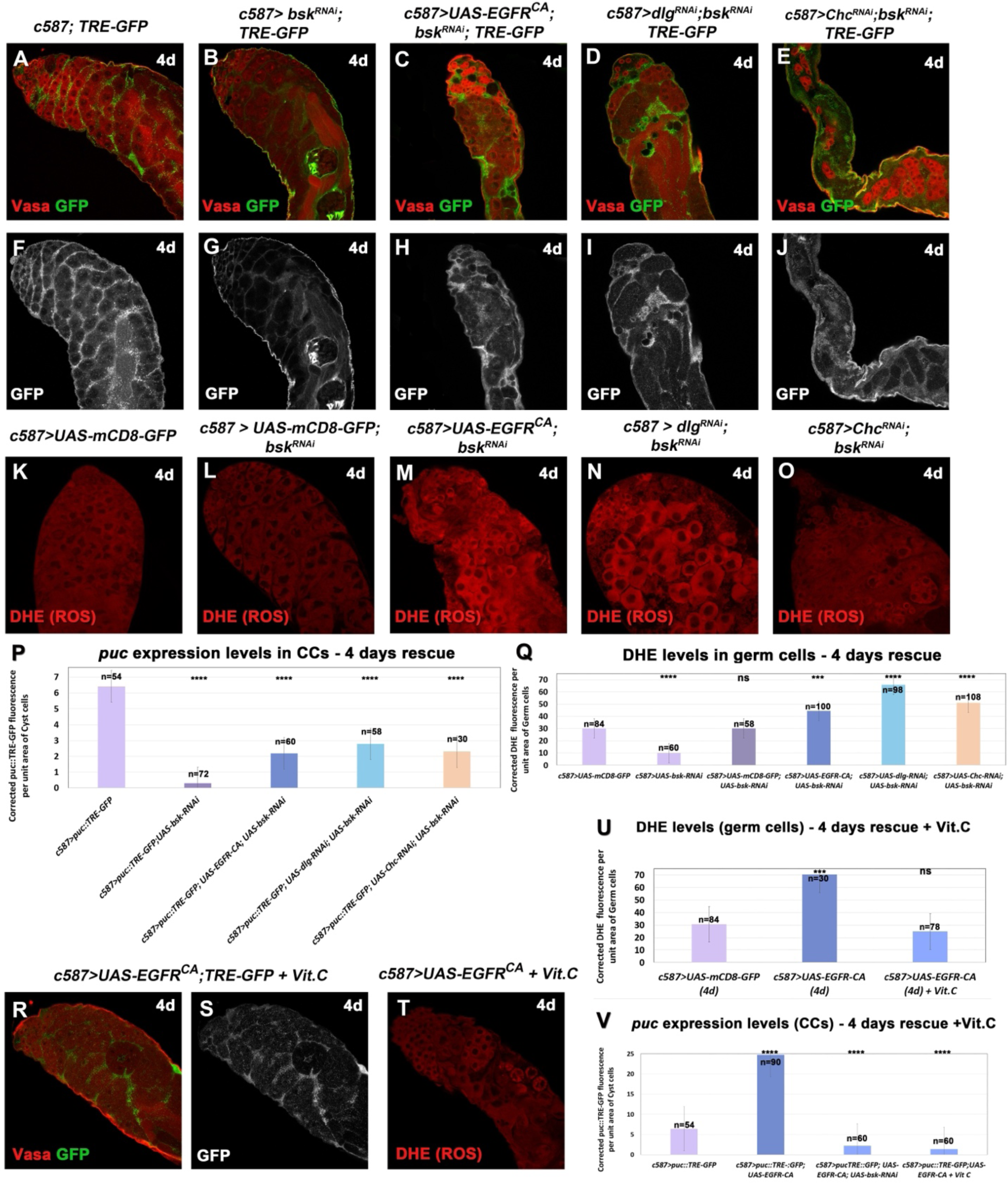
Knocking down the JUN kinase *bsk* in the background of EGFR overactivation in cyst cells, lowers JNK signalling levels in cyst cells and ROS levels in the germline. Adult testes of the indicated genotypes in the *Gal80^ts^* background. **(A-J, R, S)** *TRE-GFP* reflects expression levels of JNK reporter *puc* in CCs. (A-E) *TRE-GFP* (green; CCs); Vasa (red; germline). (F-J, S) show the *TRE-GFP* levels only (white) in directly comparable raw images. **(K-O, T)** DHE (red; germ cells) reflects ROS activation in the germ cells. Testes in (R-T) were treated with antioxidant Vitamin C (Vit.C). **(P, V)** Quantifications of corrected fluorescent *puc::TRE-GFP* levels in CCs. **(Q, U)** Quantification of corrected fluorescent ROS levels in germ cells. *UAS* activated at 30°C for 4 days (4d). Each individual sample was compared to control (error bars: standard error; ns: not significant; *p<0.05; **p<0.01; ***p<0.001; ****p<0.0001). Numbers (n) in each column represent sample size. Image frames (A-O, R-T): 225μm

As next, we set to investigate the levels of ROS activation in *bsk*-rescued testes after 4 days activation (Fig.4K-O). Knocking down *bsk* in CCs (1x *UAS* line) reduced ROS levels significantly (****p<0.0001) compared to the control (*c587>UAS-mCD8*; 1x *UAS* line) (Fig.4Q; columns 2 vs 1), showing that lowering JNK levels affects basal ROS levels in the germline. Comparing *bsk*-rescued testes (containing 2x *UAS* lines) to the control and the *mCD8-bsk*-*RNAi* control (2x *UAS* lines), showed a reduction in ROS levels (Fig.4Q; columns 3-5 vs 1-2) compared to the “EGFR overactivation” testes (Fig.2N). Yet this reduction was not sufficient to bring ROS down to control (physiological) levels (as in the case of *puc* levels – Fig.4P). In order to compare DHE levels in *bsk*-rescued vs “EGFR overactivation” phenotypes (Fig. 2N vs 4Q), we normalised the control levels between the two datasets to allow the comparison (because pictures for the quantifications were taken with a different laser). Comparison of the two datasets showed that ROS levels were reduced by 30% in *bsk*-rescued testes with *EGFR^CA^* overexpression, 20% in *dlg/bsk* double-knockdowns and 43% in *Chc/bsk* double-knockdowns. Therefore, we could conclude that effective reduction of JNK levels (by knocking down *bsk)* in CCs experiencing increased levels of EGFR, not only reduced JNK levels in CCs but could reduce ROS levels in the neighbouring germ cells and prevent their death.

### Feeding flies with antioxidant Vitamin C could reduce ROS levels and germ cell death as well as JNK levels in the neighbouring cyst cells

Since our previous results have shown that reducing JNK signalling in CCs could also lower ROS levels in the germ cells and prevent their death, we wanted to know whether the opposite was also true i.e. whether reducing ROS levels in the germ cells not only prevented GCD but could also reduce JNK levels in the neighbouring CCs. To this end, we used Vitamin C (Vit.C) an antioxidant that has been proven effective in reducing levels of ROS oxidative stress previously (Senos Demarco and Jones, 2019). Flies overexpressing *EGFR^CA^* in CCs for 4 days at 30°C, were fed with Vit.C for the last 2 days prior to the dissection and the phenotypes were subsequently analysed (Fig.4R-V, 5B, 5J). The results showed that Vit.C was able to reduce ROS significantly, at levels comparable to the physiological ones (*c587* control) (Fig.4T, 4U). Importantly, feeding flies with Vit.C led to significant reduction of *puc* levels in CCs, comparable to the reduction observed when *bsk* was depleted in the same context (*bsk*-rescue) (Fig.4R, 4S and 4V-columns 3 and 4). These observations, were also combined with a partial or full rescue of the “EGFR overexpression” phenotype and GCD, that reached 70% of the testes scored after 4 days activation (Fig.5B and 5J-column 4) and 88% after 7 days activation (Fig.S4C and S4H-column 4). Interestingly, the Vit.C-rescue seemed to be more efficient than the *bsk*-rescue in the CCs (Fig.5J and S4H - columns 4 vs 3 in both).

**Figure 5:**
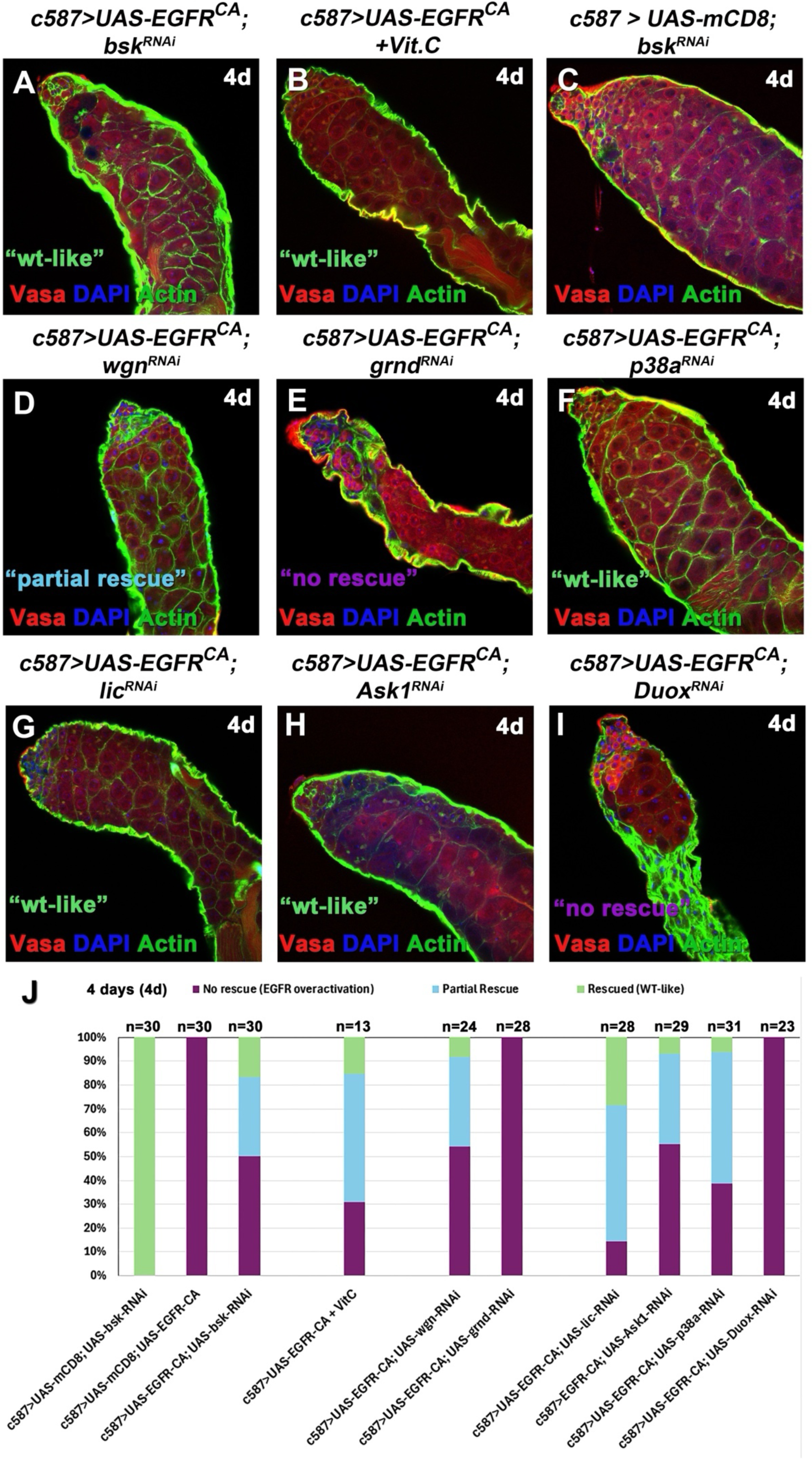
EGFR overactivation phenotypes can be partially rescued after treatment with antioxidant Vitamin C or by knocking down the JNK receptor Wengen and p38 pathway components in cyst cells. **(A-I)** Adult testes of the indicated genotypes in the *Gal80^ts^* background: Vasa (red; germline), DAPI (blue; nuclei) and Actin stained with phalloidin (here shown in the green channel; hub, CySCs, CCs and germline fusome) also in flies containing the *mCD8-GFP* transgene (since the GFP is not shown here). **(J)** Quantification of the different phenotypic classes accompanying each genotype, organized in order of phenotypic strength. *UAS* activated at 30°C for 4 days (d). Numbers (n) in each column represent sample size. Testes oriented with anterior at left. Image frames (A-I) 225μm.

### JNK receptor Wengen, Apoptosis signal-regulating kinase 1 (Ask1) and p38 signalling are involved in cyst cell-germline communication as a result of EGFR overactivation in cyst cells

In order to understand how signals from the germ cells potentially mediate JNK levels in the CCs, we looked at the two JNK TNF-receptors present in *Drosophila*, Wengen (Wgn) and Grindelwald (Grnd) (Colombani and Andersen, 2023). Knocking down *wgn* in CCs overexpressing *EGFR^CA^*, could partially rescue the EGFR overactivation phenotype (Fig.5D; column 5). Interestingly, the pattern of the three phenotypic classes (“no rescue”, “partial rescue”, “rescued”) between *wgn-* and *bsk-* rescues was very similar (Fig.5J; compare column 5 vs 3). On the other hand, knocking down *grnd* in CCs in the background of *EGFR^CA^* overexpression, could not rescue the phenotype (Fig.5E and 5J-column 6).

Apoptotic signal-regulating kinase 1 (Ask1) is a serine/threonine kinase, which senses ROS in dying and living cells, and in response to diverse stresses activates the JNK and p38 MAPK pathways (Patel et al., 2019; Santabarbara-Ruiz et al., 2019). As not only JNK but also p38 signalling is involved in apoptotic responses, we set to investigate the involvement of key components of the Ask1-p38 cascade in our system. We began our analysis by investigating the function of Ask1, Licorne (Lic), p38a and Duox in CCs. Single knockdown of any of these genes for 4 days did not have any obvious defects in differentiating germ cells or CCs (Fig. S4D-G and S4H-columns 7-10), same as already observed in the case single *bsk* depletion (Fig.5C and 5J, S4H - column 1 in both). Knockdown of *Ask1, lic* or *p38a* in CCs overexpressing *EGFR^CA^*could rescue the EGFR overactivation phenotypes, prevent GCD and restore encapsulation by the CCs, while knockdown of *Duox* could not rescue the phenotype (Fig.5F-I). Quantifying the different phenotypic classes, revealed that testes scored as partially or fully rescued represented 86% in the case of *lic*, 45% for *Ask1*, 61% for *p38a*, 0% for *Duox* (Fig.5J; columns 7-10).

In order to establish a more direct link between EGFR overactivation, GCD and the p38 pathway, the levels of phosphorylated p38 were investigated using an anti-phospho-p38 (p-p38) antibody that should recognize both activated p38a and p38b but not p38c (Patel et al., 2019). In control testes, p-p38 was clearly decorating CC nuclei but was also present in the CC cytoplasm (yellow arrowheads Fig. 6A, S5A). Immunostaining adult testes overexpressing *EGFR^CA^* or with depletion of *dlg, scrib, lgl, Chc, shi or AP-2α* in CCs, showed significant increase of p-p38 levels (****p<0,0001) (Fig.6F; columns 5-11) in the nuclei of CCs that cluster together devoid of the germ cells they normally encapsulate (yellow arrowheads Fig.6B-D, S5B-E) compared to the control (1x *UAS* or no *UAS*; Fig. 6A and 6F-columns 1 and 2). In CCs depleted of *bsk* only, p-p38 levels were significantly reduced (*p<0.05) (Fig.6E, 6F-column 4), while in the control containing two *UAS* lines (*UAS-mCD8; UAS-bsk-RNAi*) p-p38 levels were almost comparable to normal (physiological) levels (Fig. 6G and 6F-column 3 vs 1-2). p-p38 levels dropped to non-significant (ns) levels (Fig.6K; columns 6-9) in rescue experiments with double-knockdown of *bsk* with *dlg, scrib, lgl* or *Chc* (Fig. 6I, 6J, S5G, S5H), a reduction ranging between 42-78% (Fig. 6L). In the case of *EGFR^CA^*, *shi* or *AP-2α* (Fig.6H, S5I, S5J), rescued testes showed a 43-69% reduction in p-p38 levels (Fig. 6L) even though these levels were still statistically significant (***p<0.001 or ****p<0.0001) (Fig. 6K; columns 5, 10, 11) compared to the control (*c587>UAS-mCD8*). Therefore, JNK signalling regulates p38 MAPK signalling, since modifying *bsk* levels affected p38 phosphorylation and activation in CCs.

**Figure 6:**
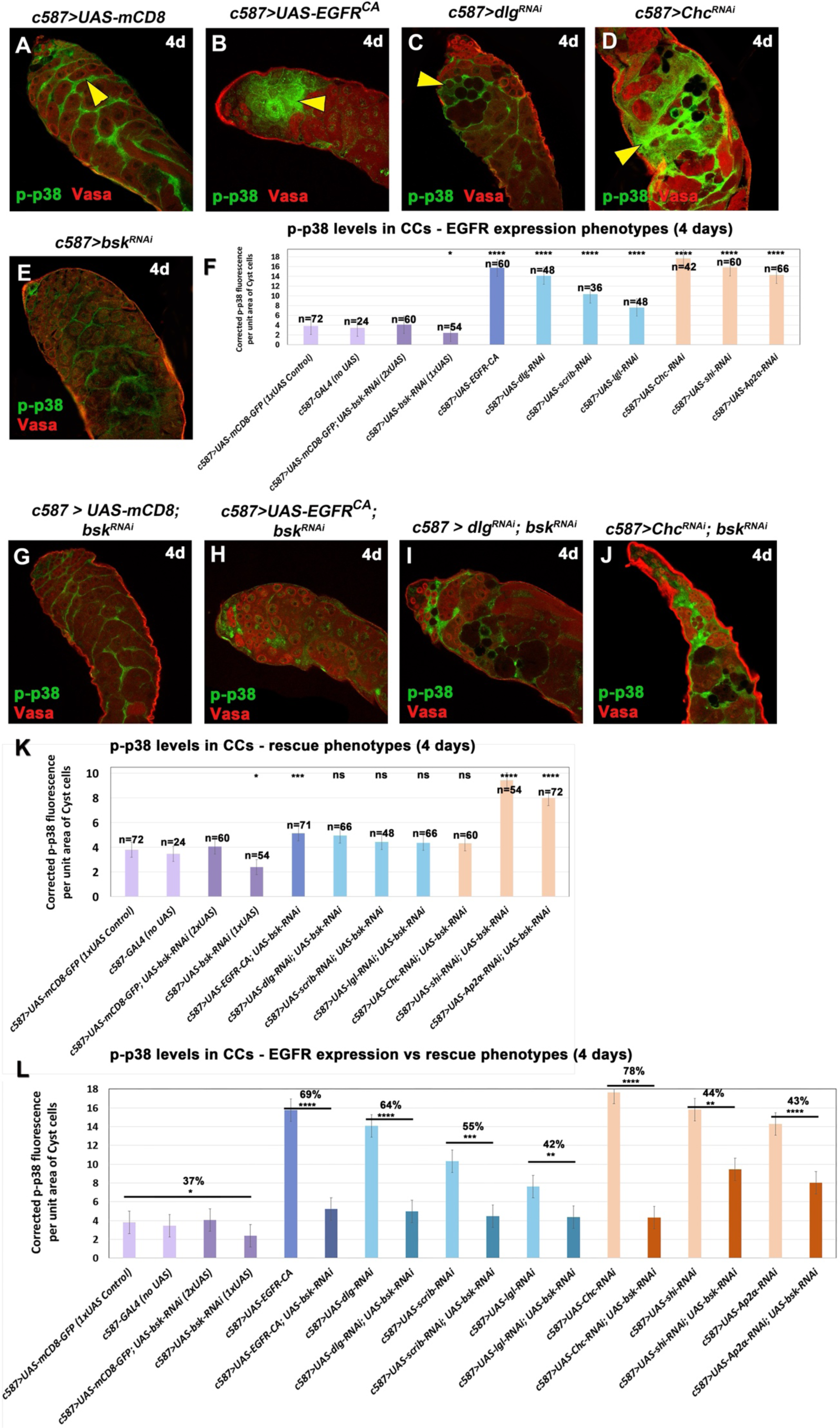
Levels of phosphorylated MAPK p38 increase in cyst cells experiencing EGFR overactivation and in response to JNK-derived cues. **(A-E, G-J)** Adult testes of the indicated genotypes in the *Gal80^ts^* background: Vasa (red; germline), DAPI (blue; nuclei) and phosphorylated p38 (p-p38) (green; CCs and nuclei), also in flies containing the *mCD8-GFP* transgene (since the GFP is not shown here). Yellow arrowheads point at CCs with high levels of p-p38. Small inset pictures show the p-p38 staining only. *UAS* activated at 30°C for 4 days (4d). **(F, K)** Quantification of corrected fluorescent p-p38 levels in CCs (and their nuclei) of indicated genotypes with “EGFR overactivation” (F) and *bsk*-rescue (K) background. Numbers (n) in each column represent sample size. For statistics in (F, K) each individual sample was compared to the *c587>UAS-mCD8* control. **(L)** Combined quantifications from (F) and (K), compare p-p38 levels in “EGFR overactivation” vs *bsk*-rescue phenotypes. For statistics, “EGFR overactivation” genotypes were compared to *bsk*-rescues, while numbers represent % of p-p38 reduction (error bars: standard error; ns: not significant; *p<0.05; **p<0.01; ***p<0.001; ****p<0.0001). Testes oriented with anterior at left. Image frames (A-Q): 225μm

### The GTPase Rab35 is required in cyst cells to regulate JNK and ROS response following EGFR overactivation

In order to investigate some of the more downstream regulators of CC-germline communication following the upregulation of the EGFR in CCs, we investigated the role of Rab35. Rab35 is a GTPase of the recycling endosome involved in plasma membrane transport, polarized trafficking, phagocytosis and exosome secretion (Bellec et al., 2018; Ochi et al., 2022; Shim et al., 2010). Use of a Rab35-GFP protein-tag in control testes, showed Rab35 localization in CCs (encapsulating spermatogonia and spermatocytes) and spermatogonia (Fig. K-K’). Rab35 was still present in CCs overexpressing *EGFR^CA^* or upon loss of *dlg* or *Chc* (Fig. M-O’). However, Rab35 was not any more visible after knocking down *Rab35* in CCs, an effect that was not associated with other visible defects in CCs and differentiating germ cells (Fig. S5L-L’).

Knocking down *Rab35* in CCs for 4 days, in the background of *EGFR^CA^* overexpression or loss of *dlg* and *Chc* (Fig.7A-E), led to a partial rescue of the EGFR overactivation phenotype, although this was less efficient compared to the *bsk*-rescues (Fig.7M: columns 4-6 vs 3 and Fig.3Q). When we compared *Rab35*-rescued vs *EGFR^CA^* overexpressing testes we observed significant reduction in *puc* expression levels in CCs (Fig.7N; column 5 vs 2) and ROS levels in germ cells (Fig.7O; column 5 vs 2). Yet, when we compared *bsk*-rescued vs *Rab35*-rescued testes reduction of *puc* levels was more significant in *bsk*-rescued testes (Fig.7N; column 5 vs 3), while reduction of ROS levels was at similar comparable levels between *bsk*- and *Rab35*-rescued testes (Fig.7O; column 3 vs 5). Thus, Rab35 emerged as one of the EGFR and JNK downstream effector genes that mediates CC-germline reciprocal communication and transduction of the death signal.

**Figure 7:**
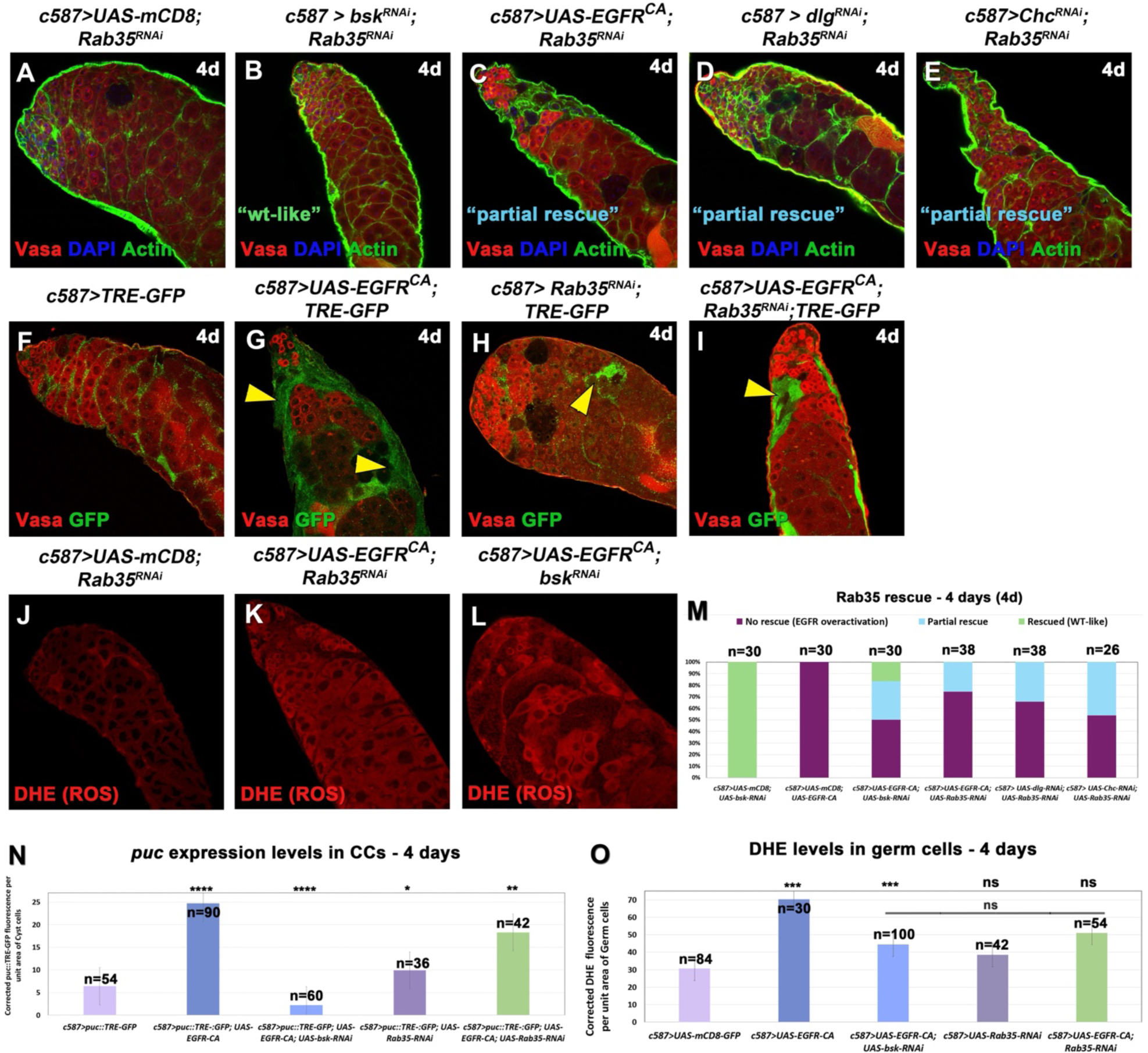
Knocking down the GTPase *Rab35* in the background of EGFR overactivation in cyst cells, lowers JNK signalling levels in cyst cells and ROS levels in the germline. **(A-L)** Adult testes of the indicated genotypes in the *Gal80^ts^* background. **(A-E)** Vasa (red; germline), DAPI (blue; nuclei) and Actin stained with phalloidin (here shown in the green channel; hub, CySCs, CCs and germline fusome) also in flies containing the *mCD8-GFP* transgene (since the GFP is not shown here). (F-I) *TRE-GFP* (green; CCs) reflects expression levels of JNK reporter *puc* in CCs; Vasa (red; germline). Yellow arrowheads: regions of increased *puc::TRE* expression. **(J-L)** DHE levels reflect ROS activation in the germ cells (red; germ cells). **(M)** Quantification of the different phenotypic classes accompanying each genotype, organized in order of phenotypic strength. **(N, O)** Quantification of corrected fluorescent *puc::TRE-GFP* levels in CCs and ROS levels in germ cells, respectively. *TRE-GFP* reflects expression levels of JNK reporter *puc* in CCs. *UAS* activated at 30°C for 4 days (d). Each individual sample was compared to control (error bars: standard error; ns: not significant; *p<0.05; **p<0.01; ***p<0.001; ****p<0.0001). Numbers (n) in each column represent sample size. Image frames (A-L): 225μm.

## Discussion

Short-range communication between closely apposed cells is critical for building functional tissues and organs and maintaining homeostasis. The *Drosophila* testis provides an excellent system to study *in vivo* how closely apposed cell types reciprocally communicate and coordinate their co-differentiation (Lim and Fuller, 2012; Papagiannouli et al., 2018; Zoller and Schulz, 2012). Our results show that the cortical polarity proteins Dlg, Scrib, Lgl and clathrin-mediated endocytosis (CME) fine-trim EGFR signalling levels and thereby influence the network of JNK, p38 and ROS signalling cross-regulation between the CCs and the differentiating (spermatogonia and spermatocyte) germ cells.

As in previous studies (Chang et al., 2019; Chang et al., 2020; Herrera et al., 2021; Herrera and Bach, 2018; Senos Demarco and Jones, 2019; Tan et al., 2017), we observe basal (physiological) levels of JNK in CySCs and CCs and ROS in the germline, although we primarily focused on the differentiating CCs and germ cells where our phenotypes are manifested. Our findings have shown that depletion of *dlg, scrib, lgl* or CME components in CCs leads to EGFR overactivation, which is accompanied by upregulation of JNK and p38 in CCs, and ROS in germ cells destined to die (Fig.8A). Germ cell death (GCD) could be reversed by acting either in the CCs through reduction in JNK signalling levels or directly in the germ cells by reducing ROS levels. More precisely, knocking down *bsk* in CCs reduced intrinsic levels of JNK and p38 as well as ROS levels in the neighbouring germ cells. Yet, a reduction of ROS levels in germ cells by feeding flies with Vitamin C, seemed to be more effective in reducing JNK levels in CCs (compared to the reverse), suggesting that oxidative stress could potentially have a more instructive role in directing CC behaviour besides triggering germline apoptosis.

**Figure 8:**
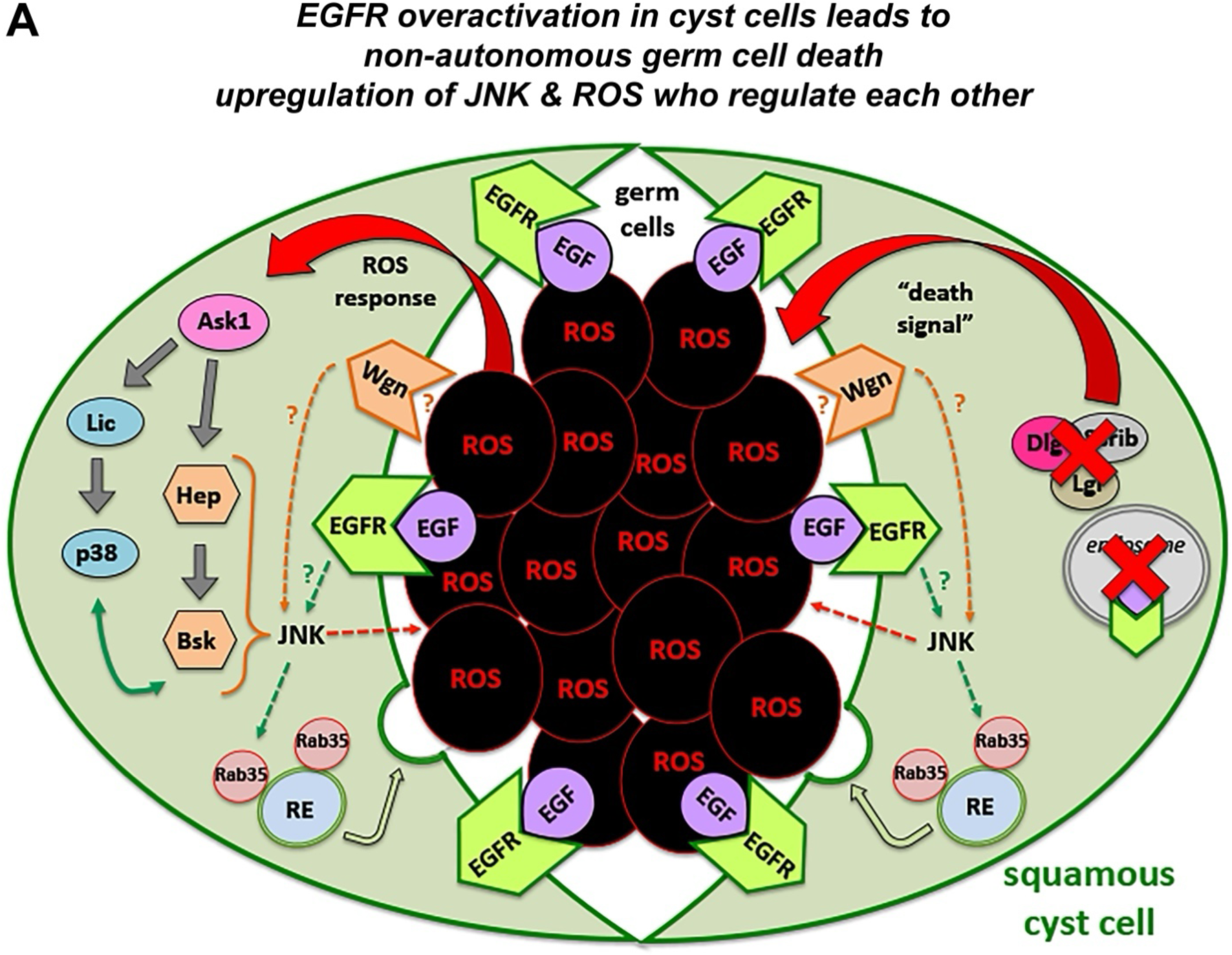
Model diagram showing the JNK, p38 and ROS interplay in cyst cell-germline cysts upon overactivation of the EGFR signalling pathway. **(A)** Loss of Dlg/Scrib/Lgl or CME components in CCs upregulates EGFR signalling that sends a “death signal” to the neighbouring germ cells (dying germ cells shown in black). This “death signal” activates ROS in the germ cells prior to their death, which could be mediated by JNK signalling, Rab35 and/or yet another unidentified molecule. ROS responds back to the CCs by sensitising Ask1 to activate the JNK and p38 MAPK signalling cascades, which signal back to the germ cells (dashed red arrow). The JNK receptor Wengen (Wgn) is also involved in activating JNK in CCs (dashed orange arrow) but whether it follows an Egr-mediated or unconventional trafficking-mediated mechanism is currently unclear (orange question mark). Rab35 as part of the recycling endosome (RE), becomes activated downstream of the JNK/p38 cascade (shown in a dashed green arrow) and contributes to the “death signal” delivery at the CC-germline inteface.

In tissues that harbour stem cell linages, JNK signalling is activated in a variety of stress responses with an instructive role in regulating the balance between homeostasis and tissue regeneration vs. degeneration, aging and death (Gan et al., 2021; La Marca and Richardson, 2020; Mundorf et al., 2019). In the *Drosophila* testes undergoing chronical stressful situations, JNK signalling seems to confer robustness by permitting Bam+/spermatogonia to dedifferentiate and renew the pool of GSCs (Herrera and Bach, 2018). Yet, stress linked to protein starvation leads to death of spermatogonia, by upregulating Spichthyn (Spict) in the encapsulating CCs (Chiang et al., 2017), which together with the lysosome promote spermatogonial phagocytic clearance. Similarly, germ cells that initiate the apoptotic pathway are eventually eliminated through the phagocytic action of JNK-upregulated CCs who extend their membranes into the dying germ cells (Zohar-Fux et al., 2022), which is very similar to our observations as well. In this study, we show a direct correlation between JNK upregulation and non-autonomous GCD, as TUNEL+ germ cells that turn on the apoptotic pathway are tightly wrapped by CCs with strong JNK-upregulation upon loss of the basal polarity Dlg/Scrib/Lgl complex and CME components. A similar mechanism of JNK-upregulation that drives GCD has been described upon loss of the apical polarity Baz/aPKC/Par6 complex in CCs, with the difference that GCD was activated only in spermatocytes and not in spermatogonia (Brantley and Fuller, 2019). However, the underlying reason that drives GCD in different germ cells in basal vs. apical components is currently unknown. A reciprocal signal, where cells with phagocytic-potential and dying cells signal at each other has been described in *Drosophila* ovaries (Serizier et al., 2022). There, the phagocytic receptor Draper in epithelial follicle cells induces non-autonomous cell death in the neighbouring nurse cells through a JNK-dependent mechanism. The MAPK p38 has been also implicated in phagocytic encapsulation of bacterial pathogens, as an immune response to better tolerate bacterial infections (Shinzawa et al., 2009). Although our data show a direct correlation between JNK and p38 activity in CCs upon EGFR overactivation, whether p38 could also contribute to the phagocytic clearance of the dying germline we observed, would require further investigation.

The EGFR signalling pathway regulates and co-ordinates a great number of fundamental functions in the *Drosophila* testis. Early studies have shown its importance in germline differentiation by directing the TA spermatogonial divisions, followed by the transition to the pre-meiotic spermatocyte program as EGFR signalling levels increase (Hudson et al., 2013; Kiger et al., 2000; Sarkar et al., 2007; Tran et al., 2000). The encapsulation of the germline by the CCs is also controlled by the EGFR, who regulates the activity of the downstream effector Rac1 that promotes the growth of CCs around the germ cells and the encapsulation strength by counteracting the function of Rho (Sarkar et al., 2007). A recent study highlighted the role of EGFR signalling in keeping the balance of CySC maintenance vs CC differentiation, through regulation of autophagy-induced lipid breakdown (Senos Demarco and Jones; Senos Demarco et al., 2020). In CySCs, EGFR stimulates autophagy through an AP-1/Fos-mediated transcription of autophagy related genes. Early CCs moving away from the hub suppress lipophagy via TOR to allow the CySC-to-CC fate switch to occur, while defective autophagy expands the CySC population in testes (Senos Demarco and Jones; Senos Demarco et al., 2020). On the other hand, ablation of CySCs activates the otherwise quiescent hub cells to turn on the EGFR signalling and resume the proliferation capacity of hub cells that can trans-differentiate and replenish CySCs (Greenspan et al., 2022). Thus, the EGFR plays an important role not only in promoting the germline differentiation and survival, but also in safeguarding the proper fate balance within the somatic lineage (hub, CySCs and CCs). Taken together, EGFR emerges as an upstream master regulator of testis spermatogenesis in normal as well as stressful conditions, with the basal polarity Dlg/Scrib/lgl complex and CME having an instructive role in EGFR-mediated functional homeostasis that is also intimately linked to JNK, p38 and ROS signalling.

ROS can activate JNK and p38 through a positive feedback loop, and thereby regulate the balance between apoptosis and autophagy vs cell competition, immunity, stem cell and tissue regeneration in many tissues and organs across species (Binh et al., 2022; Diwanji and Bergmann, 2020; Kucinski et al., 2017; Santabarbara-Ruiz et al., 2015) [reviewed also in (Herrera et al., 2021; La Marca and Richardson, 2020; Serras, 2022; Sinenko et al., 2021; Sui et al., 2014; Yue and Lopez, 2020; Zhou and Sakurai, 2022)]. Increased levels of ROS can propagate paracrine signals that are sensed by neighbouring healthy cells, which involve activation of Ask1 and subsequent phosphorylation and activation of JNK and p38 (Esteban-Collado et al., 2023; Serras, 2022). In the mammalian testes, delicate levels of ROS mediate a positive feedback loop that sustains spermatogonial stem cell self-renewal by activating the MAPK p38 (Morimoto et al., 2019). In the *Drosophila* testis, upregulation of ROS levels in the germline have already been observed, upon disruption of the antioxidant response of Keap1 (Kelch-like ECH-associated protein1)/Nrf2 (NF-E2-related factor 2), or mitochondrial fission by Drp1 (dynamin-related protein 1) (Senos Demarco and Jones, 2019; Tan et al., 2017). In both cases, increased levels of ROS result in loss of GSCs (and spermatogonia in the case of *Drp1* loss) and upregulation of the EGF-ligand Spitz, which overactivates the EGFR signalling pathway in the neighbouring CCs (Senos Demarco and Jones, 2019; Tan et al., 2017). Our findings show that elevated levels of ROS in the germline can also result by directly modifying EGFR signalling levels in the CCs, like the ones observed upon loss of the Dlg-polarity and CME components or forced expression of EGFR in CCs, all leading to death of both spermatogonia and spermatocytes (Papagiannouli et al., 2019). We also saw that upregulation of EGFR in the CCs and ROS signalling in germ cells, is linked to JNK and p38 activation in CCs. Importantly, our rescue experiments support a model of reciprocal CC-germline communication between JNK/p38 and ROS signalling that controls germ cell survival, since suppressing *bsk* reduces p38 and ROS levels, while antioxidant treatment reduces ROS and normalizes JNK levels at the same time. Only few studies so far link the Dlg/Scrib/Lgl module to ROS oxidative stress, besides neoplastic tumours (Bunker et al., 2015; La Marca and Richardson, 2020). For example, Scrib regulates ROS levels and autophagy in mouse intestinal stem and epithelial cells, which is deregulated in inflammatory bowels disease (Sun et al., 2023). Moreover, loss of function of Dlg and Rab5 endocytosis are associated with ROS activation in fly nephrocytes (the equivalent to human podocytes) that affect slit-diaphragm integrity (Xi et al., 2024). Along these lines, we favour a model where EGFR/Ras/MAPK, JNK/p38 and ROS are part of a bidirectional feedback mechanism, in which EGFR upregulation can be both the cause and/or the consequence of ROS activation in the germline, while Dlg/Scrib/Lgl and CME act as guardians of this signalling homeostasis.

Our results have also indicated the importance of the TNFα receptor Wengen (Wgn) in CCs experiencing EGFR overactivation, while Grindelwald (Grnd) seems not to be part of this signalling wiring. The fact that knockdown of *wgn* in CCs could reverse GCD (resulting from EGFR overactivation) in a rescue-pattern and efficiency that was comparable to that of *bsk*- and *Ask1*-rescue, let us think that Wgn and Ask1 could be the “messengers” of the ROS-germline-mediated signal inside the CCs to activate JNK signalling. The two *Drosophila* TNFα receptors, Wgn and Grnd, seem to have common functions albeit following a context-dependent mode of action within the cells they are functionally active. Egr/TNFα can bind Wgn with an affinity that is three times weaker compared to that of Grnd (Palmerini et al., 2021), which could explain why in our system Wgn is involved in JNK activation, as a result of ROS overactivation in germ cells. Our hypothesis is further reinforced by studies showing that Wgn favours a more “local”, close-range function (e.g. in neurons that rely on a local source of Egr) vs Grnd who seems to be involved in more systemic stress responses (Palmerini et al., 2021). Along this line, it was previously shown that initiation of reproduction in *Drosophila* males, activates the release of Egr from the smooth muscle sheath that surrounds the *Drosophila* testis to activate JNK signalling through Grnd. In this case, Grnd increases JNK in CCs at levels which are significant (*p<0.05) (as shown by *puc::TRE-GFP*) (Chang et al., 2019) but not as high as in our experimental context (****p<0.001) that is mediated by Wgn. Similarly, the Egr(smooth muscle)/Grnd(CCs) pathway is also activated upon protein starvation to promote the recovery of CySCs upon protein refeeding (Chang et al., 2020). Interestingly, Wgn has been shown to function in unconventional Egr-independent ways by becoming internalized in intracellular vesicles to regulate tracheal development (Letizia et al., 2023). Wgn has also been implicated in suppressing autophagy-dependent lipolysis in the *Drosophila* gut enterocytes to maintain homeostasis independently of Egr (Loudhaief et al., 2023), a function which in the *Drosophila* testis has been linked to TOR, who counteracts the function of EGFR/AP-1 in promoting lipophagy in early CCs (Senos Demarco and Jones, 2020; Senos Demarco et al., 2020). Therefore, we cannot exclude the possibility for Wgn to act in the *Drosophila* testis in an Egr-independent way that rescues the GCD phenotype in parallel to interfering with lipophagy.

We further showed that knockdown of the GTPase Rab35 could rescue GCD downstream of EGFR and JNK overactivation in CCs. Rab35 is a multifunctional protein of the recycling endosome, involved in membrane and endocytic trafficking that affects cytokinesis, cell adhesion, exosome release and axon elongation (Allaire et al., 2013; Klinkert and Echard, 2016; Kouranti et al., 2006; Maejima et al., 2023). Rab35 controls several aspects of apicobasal polarity with subcellular precision by regulating actin and microtubule dynamics as well as membrane PtdIns(4,5)P2 homeostasis (Francis et al., 2022; Kouranti et al., 2006; Ochi et al., 2022). By affecting actin and microtubule remodelling, Rab35 has been shown to mediate the transport of Cdc42 and Rac1 to the plasma membranes of filopodia-like protrusions involved in phagocytosis in *Drosophila*. Loss of Rab35 could rescue spermatocyte GCD when the apical Baz/aPKC/Par6 complex is disrupted in *Drosophila* testis, in line with our observations when the basolateral Dlg/Scrib/Lgl complex is depleted in CCs (Brantley and Fuller, 2019). Although it is not clear how Rab35 is activated in our system downstream of EGFR and JNK signalling pathways, we hypothesize that Rab35 has a key role in delivering the death signal from the CCs to the germline via an exocytosis or phagocytosis related mechanism or alternatively by membrane-targeting of critical components (or receptors) to the CC-germline interface.

This study highlights a bidirectional feedback mechanism that underlies CC-germline reciprocal communication that controls the signalling strength among EGFR, JNK, p38 and ROS within the *Drosophila* testis cysts. Cortical polarity components Dlg/Scrib/Lgl and Clathrin-mediated endocytosis (CME) play a pivotal role in maintaining the signalling output and physiological equilibrium that enables germline differentiation and spermatogenesis to proceed without problems. Even more, this work provides an important example on how polarity components cooperate with cellular trafficking and signalling mechanisms cell-intrinsically and across cells to build functional tissues and organs. It also provides a paradigm of how tissues can employ existing cellular networks and switch their physiological function to one that promotes cell death upon a certain threshold, when regeneration or alleviating the side-effects of stress responses has failed and cannot no longer be reversed.

## Material and Methods

### Fly stocks and husbandry

The following stocks were from the Bloomington Stock Center (BL) Indiana: *UAS-scrib-RNAi^TRiP.HMS01490^*, *UAS-AP2α (adaptin)-RNAi^TRiP.HMS00653^*, *UAS-shi-RNAi^TRiP.JF03133^*, *UAS-EGFR^CA^* (BL9533), *UAS-mCD8-GFP* (BL5139), *Pin/CyO; UAS-mCD8-GFP* (BL5130), *αtub84B-GAL80^ts^/TM2* (BL7017), *αtub-GAL80^ts^; TM2/TM6B,Tb* (BL7019), *UAS-Basket-RNAi^TRiP.JF01275^* (BL31323), *UAS-Rab35-RNAi^TRiP.JF02978^* (BL28342), *UAS-Ask1-RNAi^TRiP^*^.*HMS300464*^ (BL32464), *UAS-wengen-RNAi^TRiP.HMC03962^* (BL55275), *UAS-Nox-RNAi^TRiP.HMS00691^* (BL32902), *UAS-Duox-RNAi^TRiP.HMS00629^,* (BL32846). The following stocks used in this study were from the Vienna *Drosophila* RNAi Center (VDRC) Austria: *UAS-dlg-RNAi^v41134^, UAS-dlg-RNAi^v41136^/TM3, UAS-lgl-RNAi^v109604^, UAS-lgl-RNAi^v51247^, UAS-Chc-RNAi^v103383^, UAS-Chc-RNAi^v32666^, UAS-p38a-RNAi^v342386^, UAS-grindewald-RNAi^v104538^, UAS-lic-RNAi^v106822^, Rab35-FlyFos (v318284)*. All RNAi stocks used in this study have been effective in knocking down the corresponding genes in previous studies (Brantley and Fuller, 2019; Fujisawa et al., 2020; Papagiannouli et al., 2019; Patel et al., 2019).

The *c587*-*GAL4* was obtained from Margaret Fuller, JNK reporter *TRE-GFP (attP16)* thereafter mentioned as *puc::TRE-GFP* (Chatterjee and Bohmann, 2012) was obtained from David Bilder. Other fly stocks used in this study are described in FlyBase (www.flybase.org). All *UAS-gene^RNAi^* stocks are referred to in the text as *gene^RNAi^* for simplicity reasons. Knockdowns were performed using the *UAS-GAL4* system (Brand and Perrimon, 1993) by combining the *UAS-RNAi* fly lines with the cell-type specific *c587-GAL4* driver (Kai and Spradling, 2004) and *αtub-Gal80^ts^*(Lee and Luo, 1999).

For the phenotypic analysis in adult *Drosophila* testes: *c587*-GAL4; *αtub-Gal80^ts^; UAS-mCD8-GFP* or *c587*-GAL4*; αtub-Gal80^ts^*flies were crossed to *UAS-gene^RNAi^* flies. Crosses were raised at 18°C until adult flies hatched. Then adult males 1-3 days old with the correct genotype (along with few females in order to mate) were shifted at 30°C for 4 or 7 days depending on the experimental needs and the phenotypes were analysed.

### Immunofluorescence staining and microscopy

For whole mount testes immunostaining, testes were dissected in PBS, fixed for 20min in 8% formaldehyde (FA) and rinsed twice in 1% PBX (1% Triton-100x in PBS). Testes were blocked in 5% Bovine Serum Albumin in 1% PBX for 1h. Testes were incubated with primary antibodies over-night at 4°C and the following day with the secondary antibodies for 2h at room temperature (RT) in the dark (Papagiannouli et al., 2019). For testes immunostaining in the presence of GFP, 1% PBT (1% Tween-20 in PBS) was used instead of 1% PBX in all steps. For the TUNEL assay, tissue was processed as described above except that after fixation, the protocol from In Situ Cell Death Detection Kit (TMR Red, Sigma/Roche) was followed (Papagiannouli et al., 2019). For the Mmp1 staining, DHSB antibodies 3B8D12, 5H7B11, 3A6B4 where mixed in equal amounts (1:1:1 ratio) and the mix was used in a 3/10 dilution. Testes were blocked O/N and incubated in primary antibody mix for 72h. Staining with the polyclonal rabbit Phospho-p38 MAPK antibody gave unspecific sticky staining in spermatocytes. Staining was optimised by preabsorbing the antibody for 2 days with *c587>UAS-mCD8-GFP* testes, before use. Testes were mounted in ProLong® Gold Antifade with DAPI (P36931; Thermo Fischer Scientific).

The monoclonal antibodies used in this study: anti-Vasa (1/10; rat), anti-Mmp1 antibodies 3B8D12, 5H7B11, 3A6B4 (each used in 1/10 dilution) were obtained from the Developmental Studies Hybridoma Bank developed under the auspices of the NICHD and maintained by The University of Iowa, Department of Biological Sciences, Iowa City, IA 52242. Polyclonal chicken anti-GFP (13970; 1/10,000) was from Abcam, polyclonal rabbit Phospho-p38 MAPK (Thr180/Tyr182) Antibody (9211; 1/200) from Cell Signaling. Filamentous (F-actin) was stained with Alexa Fluor phalloidin 546 (1/300, Thermo Fischer Scientific) and DNA with DAPI (Thermo Fischer Scientific) or in DAPI containing mounting medium. Following secondary antibodies were used: donkey anti-mouse Alexa Fluor-546 and Alexa Fluor-647, donkey anti-rat Alexa Fluor-647, donkey anti-rabbit Alexa Fluor-488 and donkey anti-chicken Alexa Fluor-488 from Thermo Fischer Scientific (1/500).

Treatment of flies with antioxidant Vitamin C was performed by feeding male flies with food containing 100mM Vitamin C (L-ascorbic acid; Merck – A4403) for the two days before dissection.

Confocal images were obtained using a Zeiss LSM880 with Airy scan (1024×1024px, 225μm image frame) (University of Greenwich at Medway Campus). Pictures were finally processed with Adobe Photoshop 7.0.

### Quantification of fluorescent images and Statistical analysis

Quantifications were done using FiJi/ImageJ by measuring “Corrected Total Cell Fluorescence” (CTCF) [CTCF = Integrated Density - (Area of elected cell X Mean fluorescence of background readings)] using Excel. Statistical analysis was performed in GraphPad/Prism. The control (in most cases *c587-GAL4* or *c587>UAS-mCD8* testes reflecting physiological levels) was compared to each individual sample using the two-sampled Mann–Whitney test (also known as Wilcoxon test). Comparisons with a P-value ≥0.05 were marked as ‘ns’ (not significant); *p<0.05; **p<0.01; ***p<0.001; ****p<0.0001.

### ROS staining protocol

A Dihydroethidium (DHE) (D11347, 10×1mg, Invitrogen™) 30mM stock solution was prepared using anhydrous DMSO (Sigma-Aldrich, cat. no. 276855) and kept in small aliquots at −20°C until used. Testes were dissected in DMEM or Schneiders medium (both worked equally well in our hands). 30mM stock was diluted to a 30μM (1/1000) working concentration with DMEM medium and mixed well with vortex, giving rise to a pink solution. Testes were incubated in this mix for 7min in the dark, on an orbital shaker at RT. DHE solution was removed and traces were removed by incubating testes 3 times on DMEM medium for 5 min. Testes were fixed for 4min in 8% formaldehyde, rinsed once with PBS, mounted in ProLong® Gold Antifade with DAPI and imaged immediately with confocal microscope.

## Author Contributions

M.A. co-designed, performed and interpreted experiments, assisted with writing the paper and supported the study with her 3MT award. F.P. designed, performed, interpreted experiments, wrote the paper and obtained funding to support the study.

## Acknowledgements

We thank the generous *Drosophila* community for fly stocks and antibodies, in particular David Bilder, Margaret Fuller, DSHB, VDRC and BDSC Centers. We are thankful to Parthive Patel for sharing reagents on the p38 pathway and valuable feedback on our work. This work was supported by Medway School of Pharmacy funding to F.P. and PhD fellowship to M.A., The Royal Society Grant (RGS\R1\221141) and University of Greenwich (Faculty of Engineering and Science) ECR-Seed Funding to F.P., Universities of Greenwich and Kent 3MT funding to M.A.. We thank Dr. Simon Richardson for access to the confocal microscope and Dr Susan Shorter for technical support.

## Notes

### Competing Interest Statement

The authors have declared no competing interest.

