## Supplementary Figures for "Interplay of EGFR, JNK and ROS signalling in soma-germline communication in the *Drosophila* testis"

### Supplemental material

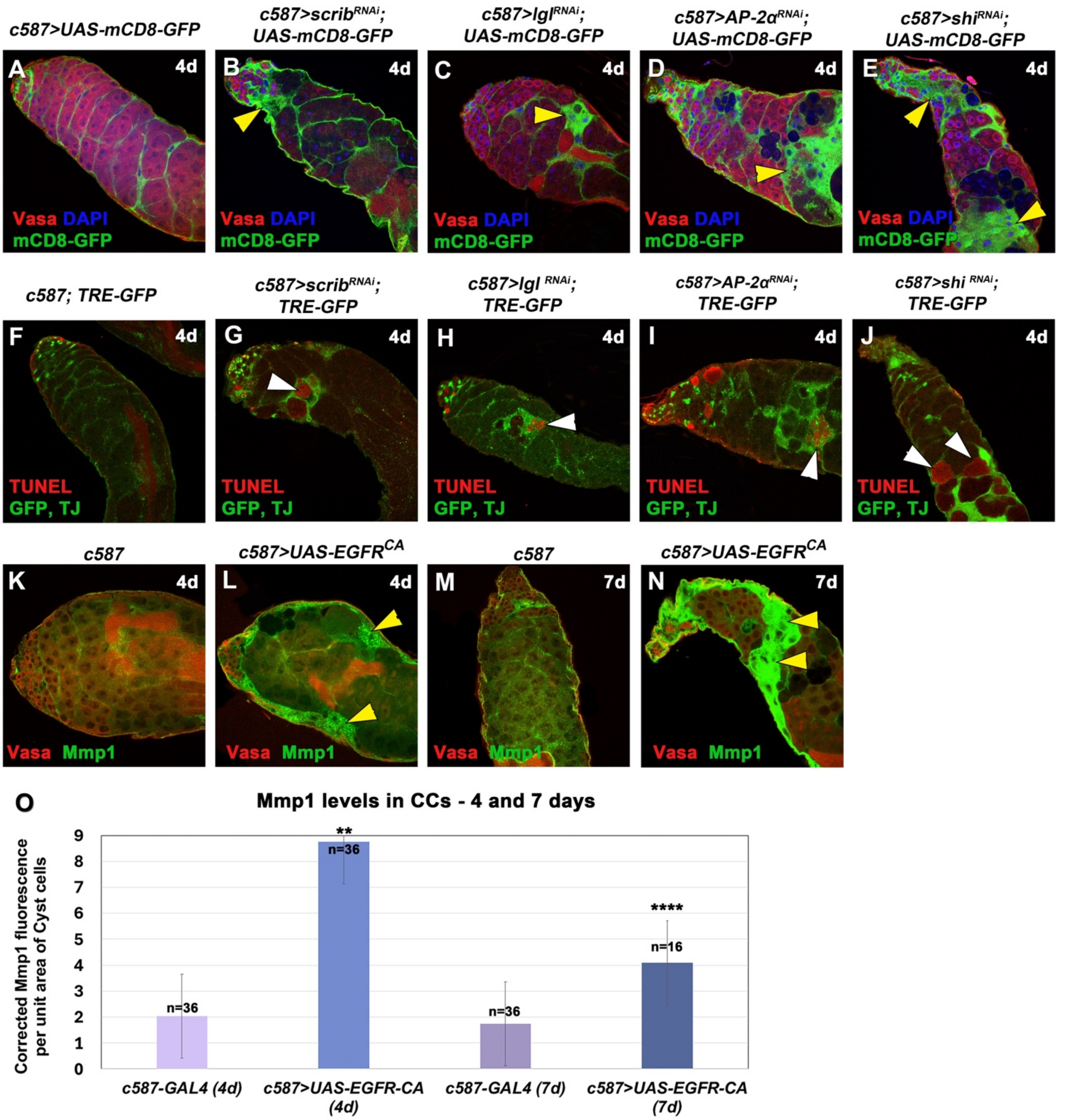


**Figure S1: Overactivation of EGFR, also via knockdown of *scrib, lgl, AP-2α* or *shi* function, in cyst cells leads to activation of JNK signaling in cyst cells and apoptosis in the neighbouring germline.** Adult testes of the indicated genotypes in the $Gal80^{ts}$ background: **(A-E)** *mCD8-GFP* (green; CCs), Vasa (red; germline), DAPI (blue; nuclei). Yellow arrowheads: mCD8+ CC regions. **(F-J)** TUNEL (red; apoptotic double-strand breaks), AP-1 responsive TRE elements corresponding to JNK reporter *puc* expression levels (*TRE-GFP*) and TJ (early CCs) (green). White arrowheads: dying germ cells (spermatogonia and spermatocytes) surrounded by CCs with upregulated JNK levels. **(K-N)** levels of the Mmp1 protein in CCs (green); Vasa (red; germline). Yellow arrowheads: Mmp1 in

CCs. *UAS* activated at 30$^{o}$C for 4 or 7 days (d). **(O)** Quantification of corrected fluorescent Mmp1 levels in CCs (4- and 7-days activation). Each individual sample was compared to control (error bars: standard error; ns: not significant; *p<0.05; **p<0.01; ***p<0.001; ****p<0.0001). Numbers (n) in each column represent sample size. Image frames (A-N): 225μm

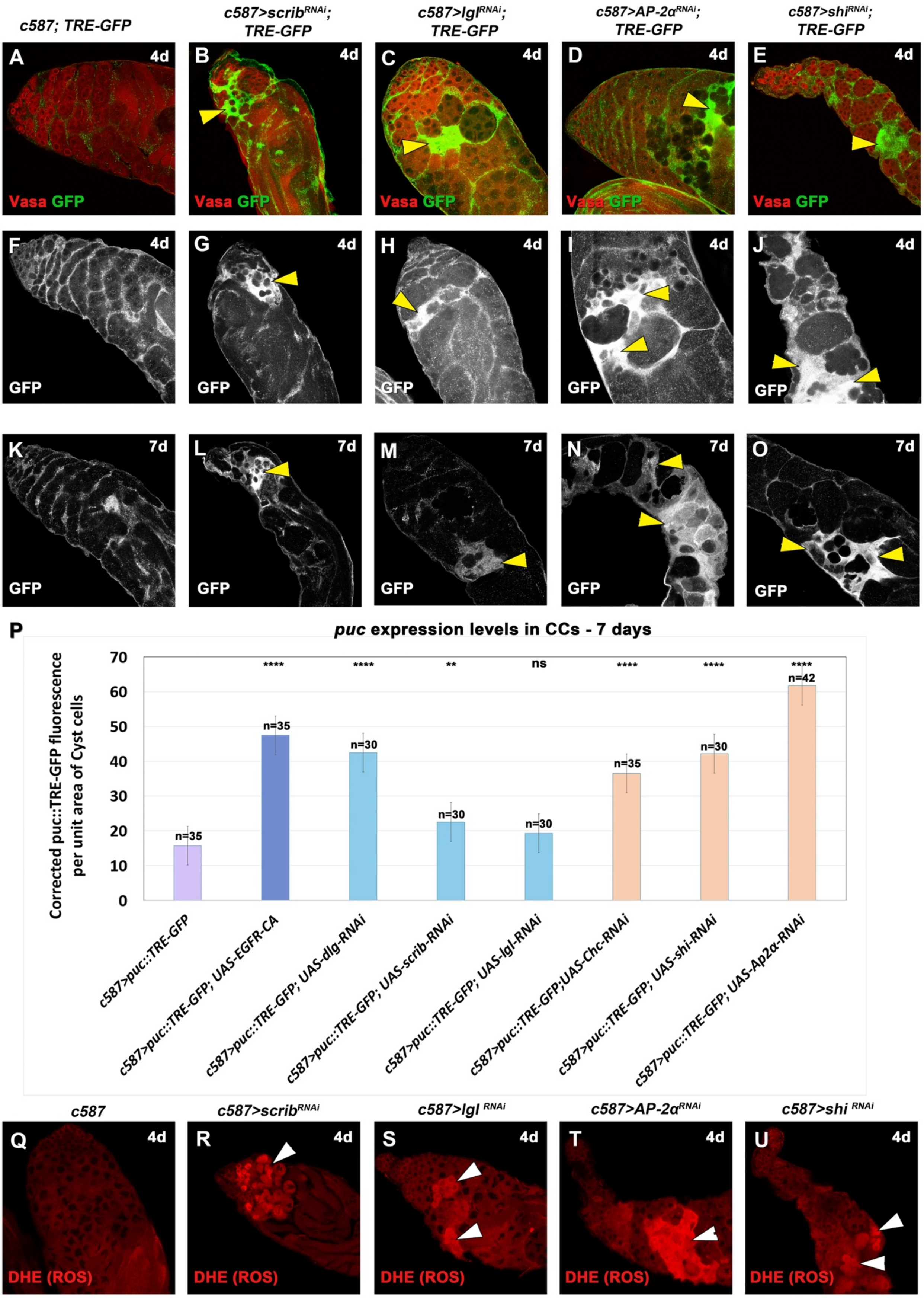


**Figure S2: Overexpression of EGFR or knockdown of *scrib, lgl* or *AP-2α, shi* function, in cyst cells leads to increased levels of JNK signaling in the cyst cells and ROS in the germline.**

Adult testes of the indicated genotypes in the *Gal80*$^{ts}$ background. **(A-O)** *TRE-GFP* reflects expression levels of JNK reporter *puc* in CCs for 4 days (A-J) and 7 days (K-O) activation. (A-E) *TRE-GFP* (green; CCs); Vasa (red; germline). (F-O) show the *puc::TRE-GFP* levels only (white) in directly comparable raw images. Yellow arrowheads: regions of *puc::TRE* overactivation in CCs. **(P)** Quantification of corrected fluorescent *puc::TRE-GFP* levels in CCs (7 days activation). Each individual sample was compared to control (error bars: standard error; ns: not significant; *p<0.05; **p<0.01; ***p<0.001; ****p<0.0001). Numbers (n) in each column represent sample size. **(Q-U)** DHE (red; germ cells) reflects ROS activation in the germ cells. White arrowheads: representative areas of ROS activation in the germline. *UAS* activated at 30°C for 4 days (4d). Image frames (A-O, Q-U): 225μm

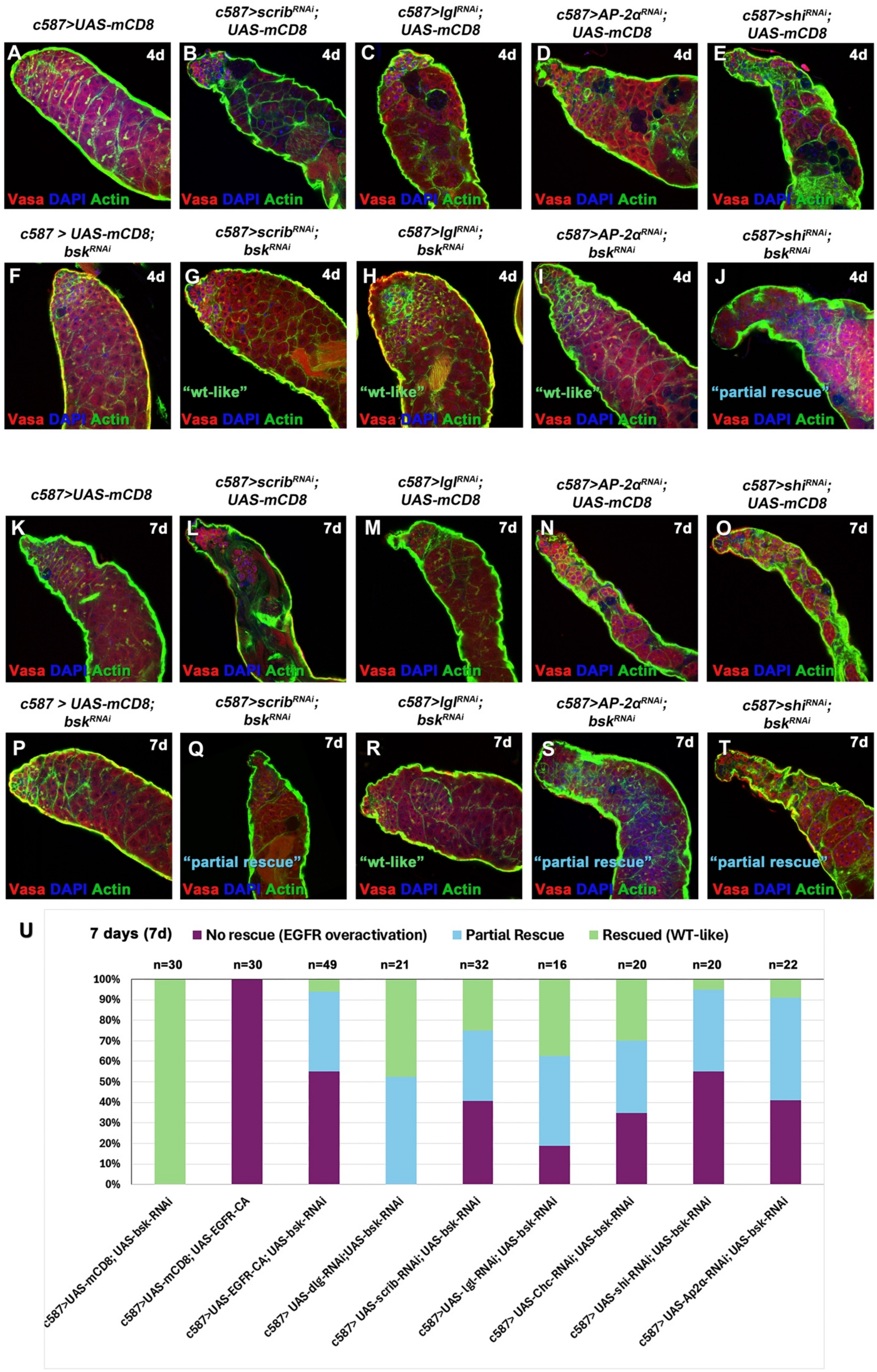
c587>UAS-mCD8
c587>scribRNAi; UAS-mCD8
c587>lglRNAi; UAS-mCD8
c587>AP-2αRNAi; UAS-mCD8
c587>shiRNAi; UAS-mCD8
A
B
C
D
E
4d
Vasa DAPI Actin
c587 > UAS-mCD8; bskRNAi
c587>scribRNAi; bskRNAi
c587>lglRNAi; bskRNAi
c587>AP-2αRNAi; bskRNAi
c587>shiRNAi; bskRNAi
F
G
H
I
J
"wt-like"
"partial rescue"
K
L
M
N
O
7d
P
Q
R
S
T
U
7 days (7d)
No rescue (EGFR overactivation)
Partial Rescue
Rescued (WT-like)
n=30
n=30
n=49
n=21
n=32
n=16
n=20
n=20
n=22
100%
90%
80%
70%
60%
50%
40%
30%
20%
10%
0%
c587>UAS-mCD8; UAS-bsk-RNAi
c587>UAS-mCD8; UAS-EGFR-CA
c587>UAS-EGFR-CA; UAS-bsk-RNAi
c587> UAS-dlg-RNAi;UAS-bsk-RNAi
c587> UAS-scrib-RNAi; UAS-bsk-RNAi
c587>UAS-lgl-RNAi; UAS-bsk-RNAi
c587> UAS-Chc-RNAi; UAS-bsk-RNAi
c587> UAS-shi-RNAi; UAS-bsk-RNAi
c587> UAS-Ap2α-RNAi; UAS-bsk-RNAi

**Figure S3: Knocking down the JUN kinase *bsk* in cyst cells, can partially rescue the germ cell death phenotype observed upon EGFR overactivation. (A-T)** Adult testes of the indicated genotypes in the *Gal80*$^{ts}$ background: Vasa (red; germline), DAPI (blue; nuclei) and Actin stained with phalloidin (here shown in the green channel; hub, CySCs, CCs and germline fusome) also in flies containing the *mCD8-GFP* transgene (since the GFP is not shown here). *UAS* activated at 30°C for 4 (A-J) and 7 (K-T) days (d). **(U)** Quantifications of the different phenotypic classes accompanying each genotype, organized in order of phenotypic strength (4- and 7- days activation). Numbers (n) in each column represent sample size. Testes oriented with anterior at left. Image frames (A-T): 225μm.

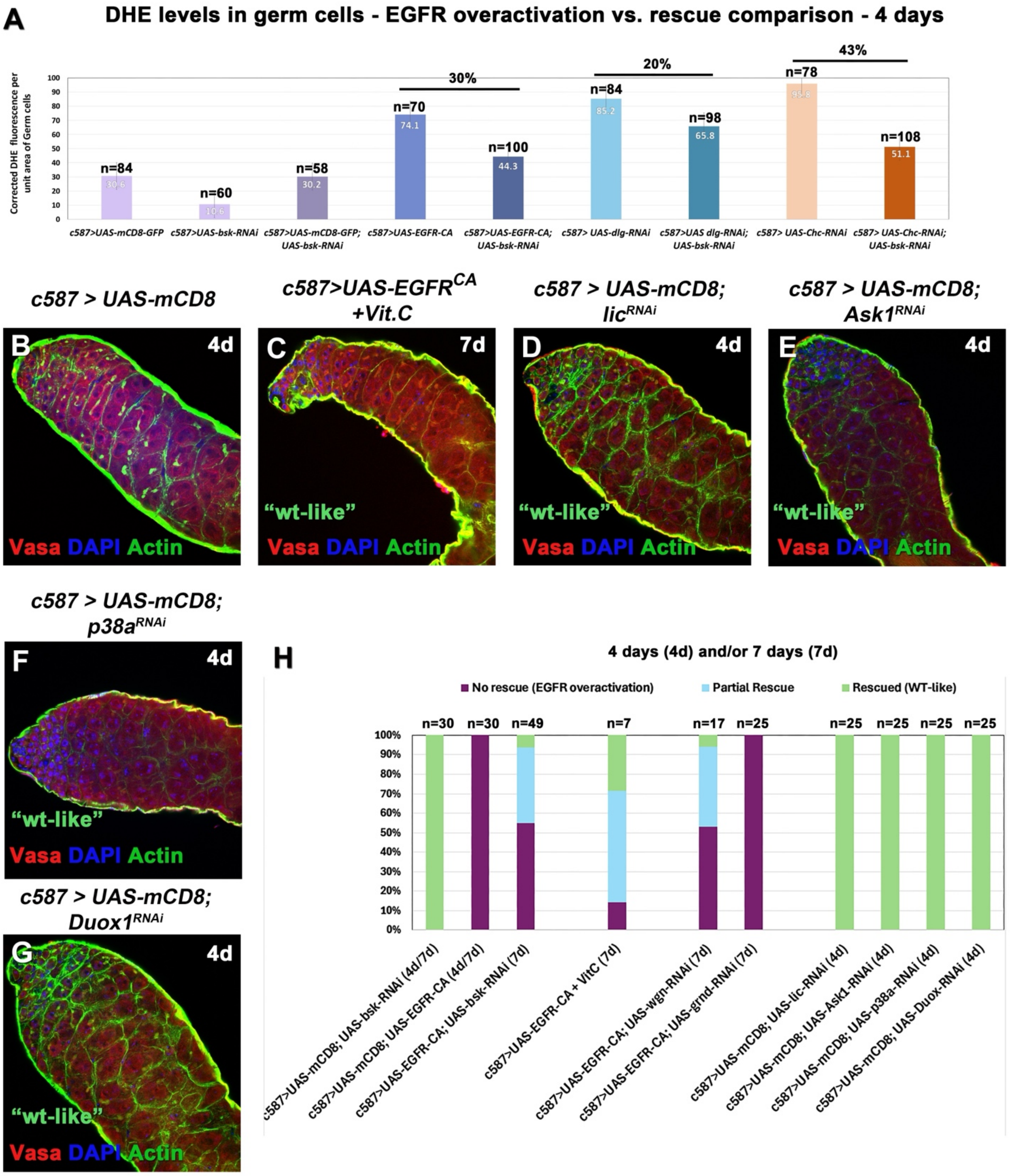


**Figure S4: (A) Double knockdown of the JUN kinase *bsk* with *scrib, lgl, Ap-2α* or *shi* in cyst cells, lowers ROS levels in the germline.** Combined quantifications of ROS levels from Fig.2N and Fig.4Q, comparing DHE levels in "EGFR overactivation" vs *bsk*-rescue backgrounds. Numbers inside the columns represent DHE levels in "EGFR overactivation" genotypes vs. rescues and % reflects this difference for each pair. Numbers (n) above each column represent sample size. As original data in Fig.2N and Fig.4Q, were obtained with a different laser, they were normalised against the *c587* controls to allow the comparison. **(B-G) Treatment with Vitamin C and knockdown of p38 pathway components in cyst cells.** Adult testes of the indicated genotypes in the $Gal80^{ts}$

background. Vasa (red; germline), DAPI (blue; nuclei) and Actin stained with phalloidin (here shown in the green channel; hub, CySCs, CCs and germline fusome) also in flies containing the *mCD8-GFP* transgene (since the GFP is not shown here). *UAS* activated at 30°C for 4 and 7 days (d). **(H)** Quantifications of the different phenotypic classes accompanying each genotype, organized in order of phenotypic strength (4- or 7- days activation) **(Matches Figure 5J).** Each individual sample was compared to control (error bars: standard error; ns: not significant; *$p<0.05$; **$p<0.01$; ***$p<0.001$; ****$p<0.0001$). Numbers (n) in each column represent sample size. Testes oriented with anterior at left. Image frames (A-G, I-L): 225μm

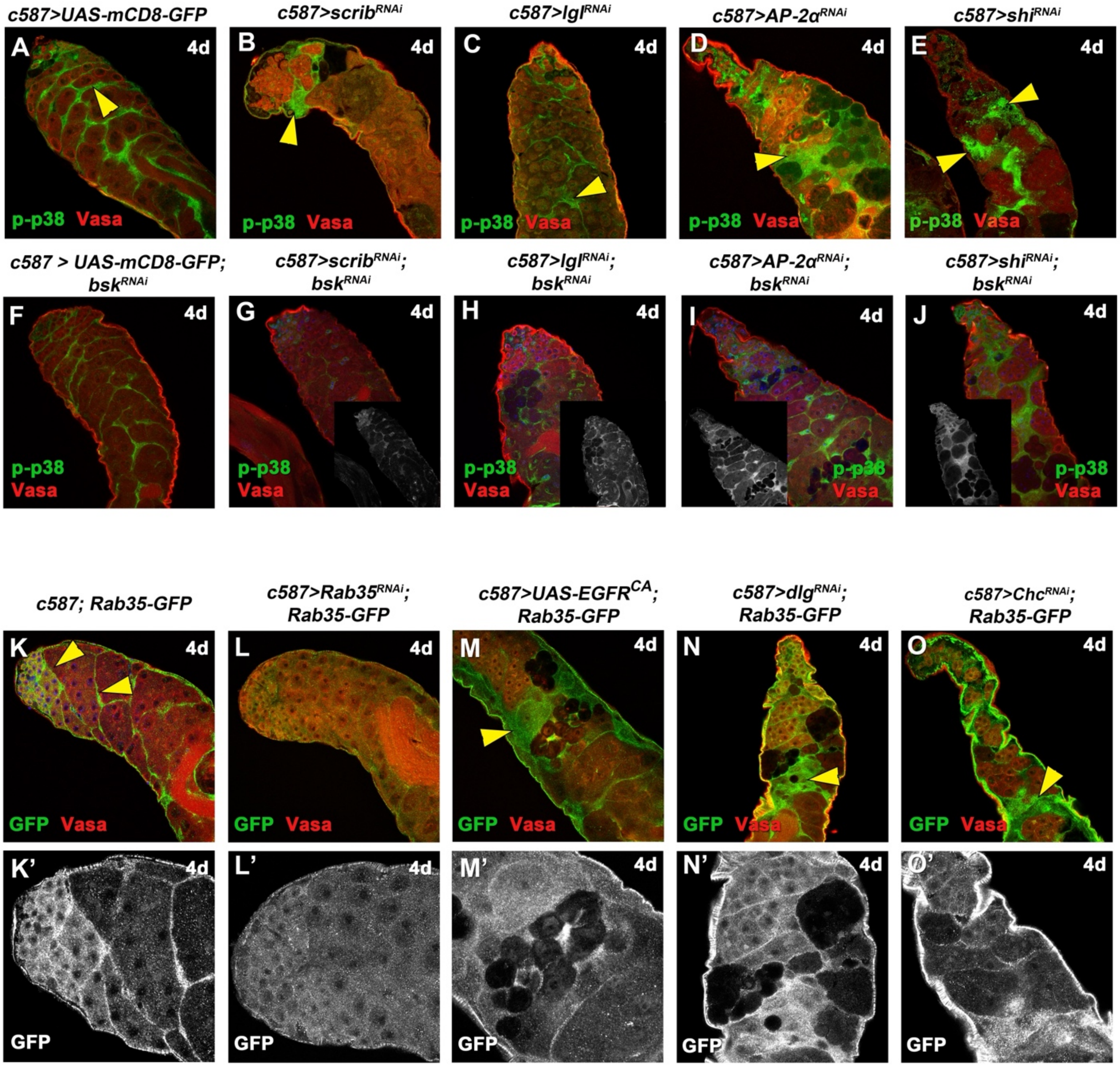


**Figure S5: (A-J) Levels of phosphorylated MAPK p38 in EGFR overactivation and rescue phenotypes after knocking down the JUN kinase *basket* in cyst cells.** Adult testes of the indicated genotypes in the $Gal80^{ts}$ background: Vasa (red; germline), DAPI (blue; nuclei) and phosphorylated p38 (p-p38) (green; CCs). Yellow arrowheads point at CCs with high levels of p-p38. Small inset pictures show the p-p38 staining only.

**(K-O’) Localization of the Rab35 GTPase in cyst cells and spermatogonia.** Adult testes of the indicated genotypes in the $Gal80^{ts}$ background. **(K-O)** Vasa (red; germline), Rab35-GFP (green; CCs and early spermatogonia). Yellow arrowheads point at Rab35 staining in the CCs. **(K’-O’)** show the Rab35-GFP staining only. **(L, L’)** shows loss of Rab35 staining after knockdown of Rab35 in CCs. *UAS* activated at 30°C for 4 days (4d). Testes oriented with anterior at left. Image frames (A-O) 225µm and (K’-O’) 112.5µm
